# Scribble and Discs-large direct adherens junction positioning and supermolecular assembly to establish apical-basal polarity

**DOI:** 10.1101/654509

**Authors:** Teresa T. Bonello, Mark Peifer

**Author notes:** **Abbreviations used:** AJ, adherens junction; Arm, Armadillo; Baz, Bazooka; BJ, basal junction; Cno, Canoe; DE-cad, *Drosophila* E-cadherin; Dlg, Discs Large; GEF, guanine nucleotide exchange factor; GFP, green fluorescent protein; Lgl, Lethal giant larvae; LRR, leucine rich repeats; MIP, maximum intensity projection; Nrt, Neurotactin; PDZ domain, post synaptic density, disc large, and zonula occludens-1 domain; SAJ, spot adherens junction; SJ, septate junctions; shRNA, short hairpin RNA; Scrib, Scribble; TCJ, tricellular junctions; WT, wildtype.

## Abstract

Apical-basal polarity is a fundamental property of animal tissues. The *Drosophila* embryo provides an outstanding model for defining mechanisms that initiate and maintain polarity. Polarity is initiated during cellularization, when cell-cell adherens junctions are positioned at the future boundary of apical and basolateral domains. Polarity maintenance then involves complementary and antagonistic interplay between apical and basal polarity complexes. The Scribble/Dlg module is well-known for promoting basolateral identity during polarity maintenance. Here we report a surprising role for the Scribble/Dlg module in polarity initiation, placing it at the top of the network that positions adherens junctions. Scribble and Dlg are enriched in nascent adherens junctions and are essential for adherens junction positioning and supermolecular assembly. They also play a role in basal junction assembly. We test hypotheses for the underlying mechanisms. Our data suggest that the Scribble/Dlg module plays multiple roles in polarity initiation, via Par-1-dependent and independent mechanisms. Different domains of Scribble contribute to these distinct roles. Together these data reveal novel roles for Scribble/Dlg as master scaffolds regulating the assembly of distinct junctional complexes at different times and places.

## Introduction

Cell polarity is a fundamental property of cells, from yeast to neurons. Epithelial apical-basal polarity provides an example (Campanale et al., 2017). Precise positioning of polarity and junctional proteins at the plasma membrane allows cells to create domains with distinct biochemical properties, allowing, for example, intestinal cells to position glucose importers apically and glucose exporters basally. Cell-cell adherens junctions (AJs) reside at the boundary between apical and basolateral domains, and are key polarity landmarks. Once established, mutually exclusive apical and basal domains are maintained and elaborated by recruitment or antagonism between apical and basolateral polarity complexes, with remarkable conservation of function across species, though sometimes acting in different combinations.

*Drosophila* embryos provide a superb model for defining the mechanistic basis of apical-basal polarity establishment and maintenance (Harris, 2012). Development begins with 13 rounds of nuclear division without cytokinesis. The last four occur at the egg cortex, which provides an important polarity landmark that polarizes these mitotic divisions. Plasma membrane then moves down around each nucleus, creating ∼6000 cells. During cellularization, cell polarity is initially established, with cadherin-catenin complexes assembled into spot AJs (SAJs) at the boundary of what will become apical and basolateral domains. AJ proteins are not essential for syncytial divisions (Grevengoed et al., 2003), but in their absence cell adhesion and polarity are lost when gastrulation begins (Cox et al., 1996; Müller and Wieschaus, 1996; Sarpal et al., 2012; Tepass et al., 1996).

Thus focus turned to defining mechanisms required to properly position and assemble AJs. The polarity regulator Bazooka (Baz=fly Par3) co-localizes with cadherin-catenin complexes and is required for their apical positioning and supermolecular assembly (Harris and Peifer, 2004; Müller and Wieschaus, 1996). An apical actin-based scaffold thought to anchor Baz and AJs apically and dynein-driven apical transport also play roles (Harris and Peifer, 2005). The actin-junction crosslinker Canoe (Cno=fly Afadin) and its regulator, the GTPase Rap1, act upstream of both Baz and AJ positioning (Bonello et al., 2018; Choi et al., 2013). Activating Rap1 apically positions Cno at nascent SAJs where Cno directs Baz and AJ positioning.

Polarity is then maintained and increasingly elaborated via recruitment or antagonism between a complex network of apical and basolateral polarity complexes (Tepass, 2012). The apical domain is initially defined by the Par complex (aPKC/Par-6) which is recruited apically in a Baz- and cdc42-dependent fashion (e.g. Bilder et al., 2003; Harris and Peifer, 2005; Harris and Peifer, 2007; Hutterer et al., 2004) and then acts in opposition to Par-1 to focus and position belt AJs (McKinley and Harris, 2012; Wang et al., 2012). The Crumbs complex is localized and promotes apical membrane identity (reviewed in Bazellieres et al., 2018). The Par complex and Crumbs antagonize apical localization of basolateral polarity proteins, while the basolateral Scribble (Scrib)/Discs-Large (Dlg)/Lethal giant larvae (Lgl) module and Yurt group act in parallel to prevent basolateral spread of apical and junctional proteins (Bilder et al., 2003; Laprise et al., 2006; Laprise et al., 2009; Tanentzapf and Tepass, 2003). Much of this is mediated by phosphorylation—aPKC phosphorylation excludes basal proteins from the apical domain (reviewed in(Hong, 2018) while Par-1 phosphorylation excludes apical proteins from the basolateral domain.

Here we focus on the Scrib-module (Stephens et al., 2018; Bonello and Peifer, 2019). Dlg and Lgl regulate tissue growth in imaginal discs, precursors of the adult epidermis (Gateff and Schneiderman, 1974; Woods et al., 1996). Scrib, which promotes embryonic epithelial integrity (Bilder and Perrimon, 2000), works with Dlg and Lgl in both epithelial polarity and growth regulation (Bilder et al., 2000). They are mutually required for one another’s localizations, but whether they form a protein complex or act in parallel remains unclear. In the mature ectoderm the Scrib-module antagonizes apical domain expansion and maintains AJs, with loss resulting in ectopic localization of apical and AJ proteins along the basolateral axis (Bilder et al., 2000; Bilder and Perrimon, 2000; Bilder et al., 2003; Tanentzapf and Tepass, 2003). Similar roles in polarity elaboration and maintenance are played by the *C. elegans* homologs (Bossinger et al., 2001; Legouis et al., 2000; McMahon et al., 2001). Interestingly, later the Scrib-module colocalizes with and is required to assemble core septate junction (SJ) proteins into functional SJs (Woods et al., 1996; Zeitler et al., 2004) which like mammalian tight junctions provide epithelial barrier function. Assembly of core SJ proteins is accompanied by enhanced immobilization on the cortex, a property that is not disrupted by Dlg loss (Oshima and Fehon, 2011). Thus, the Scrib-module is required for SJ positioning and supermolecular assembly but not for formation of the SJ core complex. Scrib and Dlg also regulate assembly/localization of other supermolecular complexes like neural synapses.

Our goal is to define mechanisms directing polarity establishment, with the ultimate objective to define the full protein network involved. While the data above suggest a simple linear pathway--Rap1GEFs->Rap1->Cno->Baz->AJs→aPKC and other proteins involved in polarity elaboration-- there are already indications things are more complex. In a linear pathway, Cno is upstream of Baz and aPKC, but their loss leads to defects in Cno supermolecular assembly. Wiring of the polarity system also varies significantly between different tissues. Here we explore roles of Scrib and Dlg. In contrast to the idea that they are primarily involved in polarity elaboration and maintenance, we find they play a critical role in polarity establishment, acting at the top of the known hierarchy.

## Results

### Scrib and Dlg already localize during cellularization and are enriched near nascent AJs

Polarity establishment begins at cellularization (stage 5), at the end of which Baz and AJs are positioned apically. At gastrulation onset (stage 6), additional polarity regulators come into play, successively elaborating polarity. In the canonical model, apical Crumbs, Yurt, and Par6/aPKC complexes and the basolateral Scrib-module and Par-1 reinforce and elaborate polarity by mutual antagonism. In the mature ectoderm, Scrib and Dlg colocalize at SJs, which are basolateral to AJs (Fig. 1A). Core SJ proteins like Coracle and Neurexin do not localize until mid-embryogenesis (Baumgartner et al., 1996; Fehon et al., 1994). While Scrib and Dlg are known to localize to the lateral membrane prior to SJ formation (Woods and Bryant, 1991), their localization during polarity establishment and the precise time course were not fully characterized.

**Figure 1.**
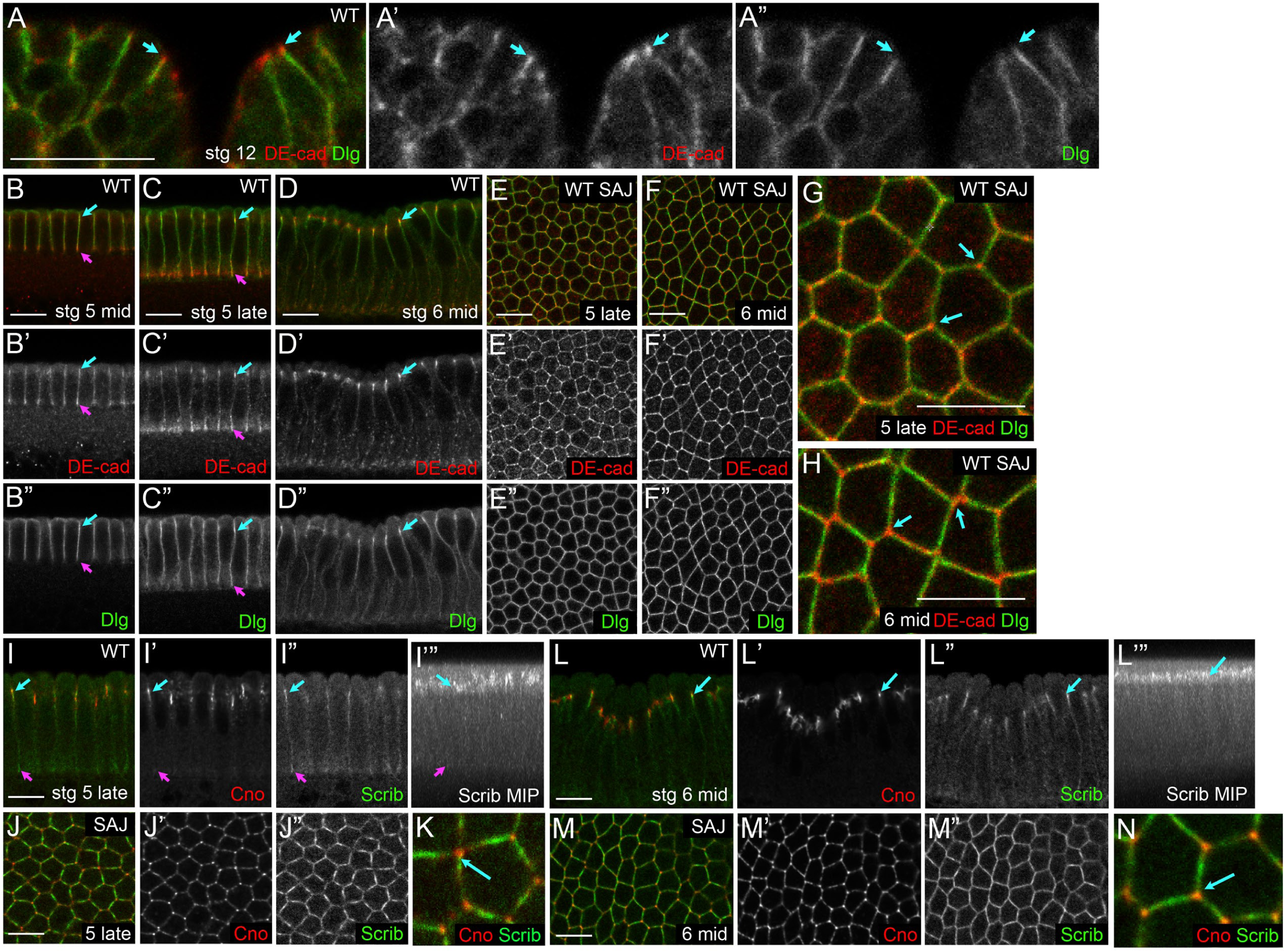
Scrib and Dlg localize to nascent AJs during cellularization. A. Stage 12 ectoderm. Dlg localizes basal to AJs (marked by DE-cad). B-N. Cellularization to early gastrulation. Dlg (B-H) and Scrib (I-N) localization relative to SAJs. B-D,I,L. Cross-sections. E-H,J,K,M,N. *En face* sections through SAJs at the level of highest enrichment. I’”,L’”. Maximum Intensity Projections (MIPs). B-C,I. Dlg and Scrib localize along the basolateral membrane during cellularization and overlap both SAJs (cyan arrows) and BJs (magenta arrows). E,J. At the level of SAJs, Dlg and Scrib localize uniformly around the cell circumference with no TCJ enrichment (G,K,cyan arrows). D,F,H,L-N. During gastrulation, Dlg and Scrib remain enriched near apical SAJs (cyan arrows). Scalebars=10µm.

If Scrib/Dlg’s roles were restricted to polarity elaboration and maintenance, one might expect they would not localize until gastrulation onset, when polarity elaboration begins, and that they would localize basolateral to AJs. However, this is not what we observed. As cellularization proceeds, AJ proteins localize to an apicolateral position in punctate SAJs (Fig. 1B,C,cyan arrows). AJ proteins also localize to smaller puncta along the lateral membrane, and are enriched in basal junctions (BJs, Fig. 1B,C,magenta arrows). In contrast to Scrib and Dlg basolateral SJ localization at stage 12, in cellularizing embryos both extended the entire length of the lateral membrane (Fig. 1B”,C”), though neither were enriched with F-actin/myosin at the furrow front. More surprising, Dlg and Scrib extended apically to overlap nascent SAJs (Fig. 1B,C,E,I,J), and by late cellularization there was clear enrichment of both Dlg (Fig. 1C”,cyan arrow) and Scrib (Fig. 1I”,cyan arrow) at the SAJ level. Maximum intensity projections (MIPs) further emphasized this (Fig. 1I’”). *en face* views revealed that while AJs proteins localize discontinuously to SAJs (Fig. 1E’, J’), Scrib and Dlg localize relatively uniformly around the circumference (Fig. 1E”,J”). While Cno and other AJ proteins are enriched at tricellular junctions (Bonello et al., 2018), Dlg (Fig. 1G,H) and Scrib (Fig. 1K,N) are not. Apical Dlg and Scrib enrichment became even more pronounced as gastrulation initiated (stage 6; Fig. 1D-D”,L-L’”,cyan arrows). These data prompted us to reconsider the possibility that Scrib/Dlg might already have roles during polarity establishment rather than only in polarity maintenance.

### Disrupting AJs does not alter Dlg apical enrichment

The close spatial relationship of SAJs, Dlg and Scrib during cellularization prompted us to investigate their functional interdependency. We first considered the hypothesis that intact AJs are necessary for apical enrichment of Dlg and Scrib. To test this hypothesis, we substantially reduced maternal and zygotic Arm (*Drosophila* beta-catenin) using RNAi. Beta-catenin is required to traffic E-cad to the plasma membrane in mammals and *Drosophila* (Chen et al., 1999; Harris and Peifer, 2004). *arm-RNAi* or mutation substantially reduces plasma membrane DE-cad (Harris and Peifer, 2004). DE-cad is lost from SAJs and BJs, and localizes to puncta aligned along the basolateral membrane and in an apical compartment above the nucleus (Fig. 2A’B’ vs C’,D’), consistent with a trafficking defect. This continues during early gastrulation (Fig. 2E’F’ vs G’,H’). However, apical Dlg enrichment was not perturbed (Fig. 2A”,B” vs C”,D”; 2E”,F” vs G”,H”) consistent with previous work showing Scrib is not lost from the plasma membrane in *arm* maternal/zygotic mutants later in development (Bilder et al., 2003). Interestingly, Cno and Baz positioning also do not require AJ function (Fig. 2I vs J; Harris and Peifer, 2004; Sawyer et al., 2009). These data suggest AJ assembly is not necessary for Scrib/Dlg apical enrichment at polarity establishment, consistent with the possibility that Scrib/Dlg act upstream.

**Figure 2.**
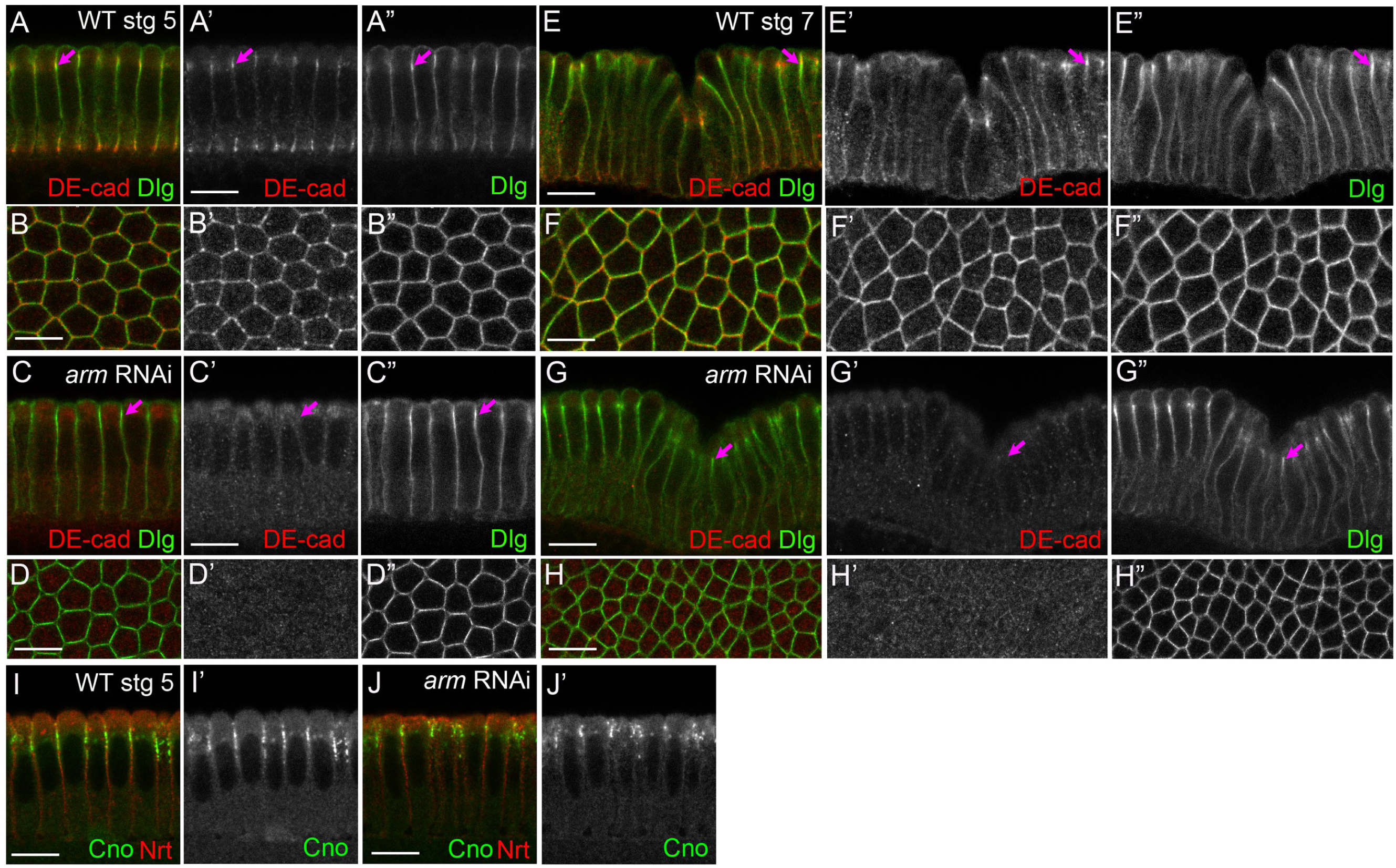
Apical Dlg enrichment does not require intact AJs. Localization of Dlg or Cno at the end of cellularization (A-D,I,J) or just prior to germband extension (E-H) in wildtype (A,B,E,F,I) or embryos expressing *arm* RNAi (C,D,G,H,J). In A-H embryos were co-stained with DE-cad to verify junctional disruption. Arrows=apical SAJs. Scalebars=10µm.

### Scrib and Dlg are essential to position AJs during polarity establishment

Polarity establishment begins with SAJ positioning and supermolecular assembly. Small cadherin-catenin clusters originating from the apical surface must be correctly positioned at the interface between apical and basolateral domains. Next, they organize into larger supermolecular assemblies (McGill et al., 2009). These are distributed around the circumference, with some enrichment at TCJs. AJ positioning requires Baz, an intact actin cytoskeleton and dynein-directed microtubule transport. The small GTPase Rap1 and its effector Cno act upstream to position and organize Baz and SAJs.

The surprising enrichment of Scrib/Dlg near nascent SAJs led us to examine whether they play roles during early polarization. To test this we maternally expressed small hairpin RNAs against *scrib* and *dlg* to reduce both maternal and zygotic transcripts. We found effective shRNAs for each, which reduced protein levels below the detection threshold of immunoblotting (Suppl.Fig. 1A,B), and replicated known cuticle phenotypes (Suppl.Fig. 2A-C). These included two *scrib* shRNAs targeting different regions of the mRNA, reducing likelihood of off-target effects.

In wildtype, cadherin-catenin assembly into apical SAJs begins early in cellularization (Fig. 3A,red arrow), and by mid-late cellularization AJs proteins are strongly enriched there (Fig. 3A’ A”,red arrows; quantified in G). This enrichment becomes clear in MIPs of multiple cross-sectional planes (Fig. 3A’”,red arrows). Cadherin-catenin complexes also accumulate in smaller puncta along the lateral membrane (Fig 3A-A’”,cyan arrows), and are enriched just apical to the furrow front in BJs (Fig. 3A-A”’,yellow arrows; Hunter and Wieschaus, 2000)). Both *scrib-RNAi* (Fig. 3B-B’”) and *dlg-RNAi* (Fig. 3C-C’”) caused pronounced AJ protein mislocalization; Arm localized to small puncta distributed nearly evenly along the apical-basal axis. Interestingly, AJ puncta not only spread basally (Fig. 3B-B’”,C-C’”, cyan arrows, D” vs E”,F”) but also mislocalized into the apical domain (Fig. 3A’” vs B’” and C’”, brackets, D vs E,F). To quantify this we measured pixel intensities along the apical-basal axis from MIPs (Fig 3G,H). This reinforced the changes in Arm localization after *scrib-RNAi*

**Figure 3.**
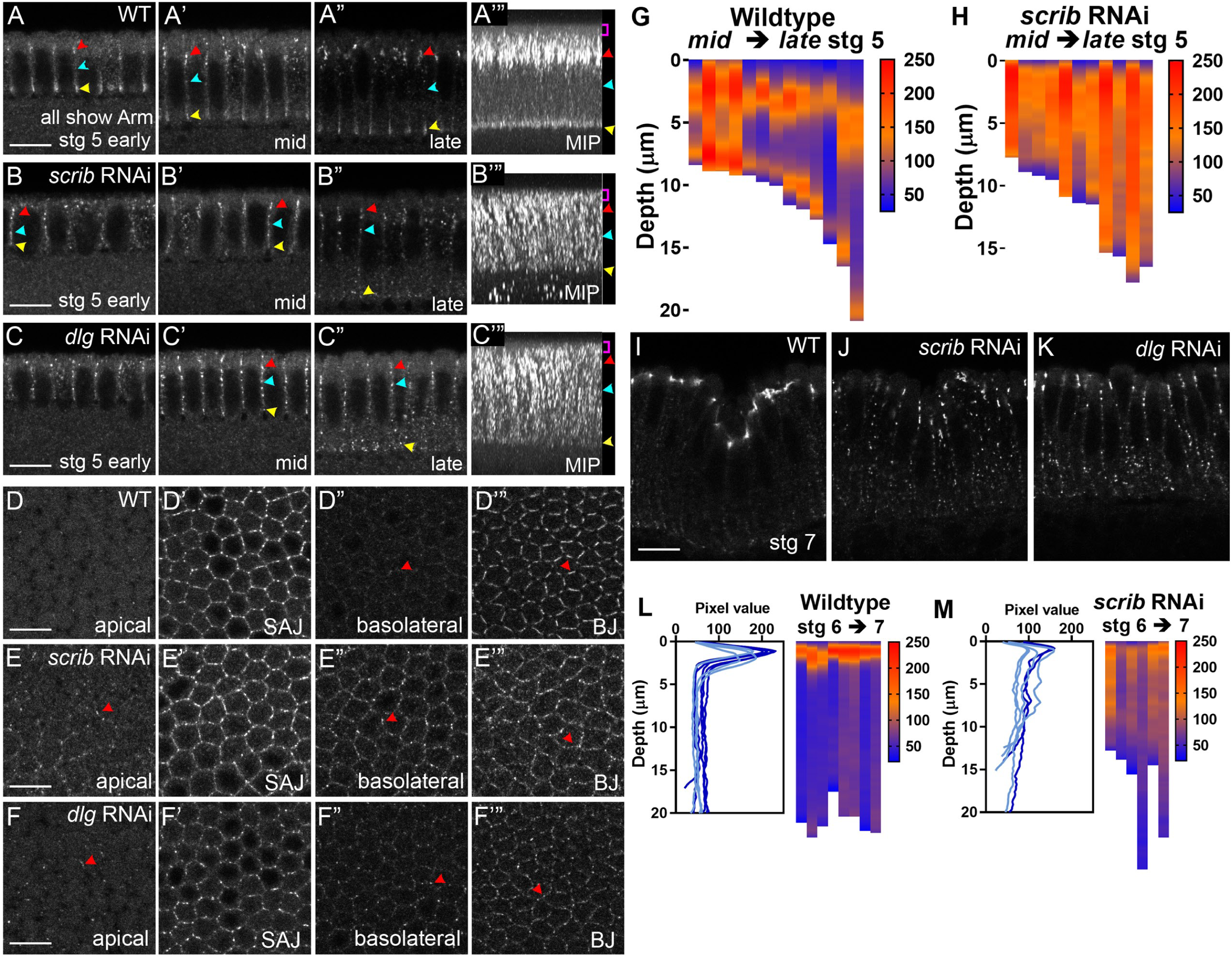
Scrib and Dlg are required for SAJ apical positioning and organization. A-C,I-K. Cross-sections. A’”-C’”. MIPs. D-F. *En face* sections through apical membrane (0µm), SAJs (−3.6µm), basolateral membrane and basal junctions (BJ). G,H,L,M. Heat maps showing Arm puncta displacement along the apical-basal axis, measured from MIPs, with pixel intensity plotted against distance from apical surface. Each vertical bar=an individual embryo. *scrib-RNAi* (B,E,J) or *dlg-RNAi* (C,F,K) disrupt Arm polarization during cellularization and gastrulation. In *scrib-RNAi* and *dlg-RNAi* embryos, Arm does not effectively enrich at apical SAJs (B,C,red arrowhead) or BJs (yellow arrowhead) and accumulates in puncta along the basolateral membrane (cyan arrowhead). G,H. Arm distribution from mid- to late-cellularization. I-K. Arm mislocalization in *scrib-RNAi* and *dlg-RNAi* embryos persists during gastrulation (stage 7). L,M. Arm distribution, stage 6-7. Scalebars=10µm.

A second junctional pool forms near the furrow tip (Fig. 3A-A’”,yellow arrows). These BJs contain core AJ proteins, but unlike SAJs do not contain Cno or Baz. BJs are regulated by the cellularization-specific protein Nullo, and disassemble at gastrulation onset (Hunter and Wieschaus, 2000). We were surprised to find that after *scrib-RNAi* or *dlg-RNAi* BJs also failed to form (Fig. 3A-A’” vs B-B’” and C-C’”,yellow arrows). Unlike punctate SAJs, BJs Arm/DE-cad form linear arrays along bicellular contacts and are excluded from TCJs (Fig. 3D’”). After *scrib-RNAi* or *dlg-RNAi*, these arrays were disrupted (Fig. 3E’”,F’”).

As gastrulation initiates, cadherin-catenin complexes in SAJs move apically and focus (Fig. 3I, quantified in L), as BJs disassemble. After *scrib-RNAi*, AJ mislocalization became even more pronounced at gastrulation onset—small cadherin-catenin complexes localized along the lateral membrane with little apical enrichment (Fig. 3J, quantified in M). Similar defects were seen after *dlg-RNAi* (Fig. 3K). Thus Scrib and Dlg play important roles in positioning and promoting supermolecular assembly of both SAJs and BJs.

### Scrib/Dlg loss disrupts Cno localization but Cno continues to co-localize with AJ proteins

Apical restriction of both Baz and AJs require Cno and its regulator Rap1. We thus examined whether Scrib/Dlg were required to apically position Cno. Cno co-localizes with Arm and Baz in SAJs from mid-late cellularization (Fig. 4A,B,brackets), and is especially enriched at TCJs (Fig. 4G’,arrows), where it forms cable-like structures (Bonello et al., 2018). Unlike core AJ proteins, Cno puncta are not found basolateral to SAJs (Fig. 4B”,red arrow) or in BJs (quantified in Fig 4I). *scrib-RNAi* disrupted this tight apical restriction during mid- (Fig. 4A vs C, brackets) and late-cellularization (Fig. 4B vs D, brackets). Cno puncta extended both apical (Fig. 4B” vs D”,yellow arrows, G vs H) and basal to the usual position of SAJs (Fig. 4B” vs D”,red arrows, G” vs H”), though remnant apical enrichment in the top half of the cortex remained (quantified in Fig. 4I vs. J). Cno organization into apical supercellular cables was lost (Fig. 4B” vs D”, brackets), and strong TCJ enrichment was reduced but not lost (Fig. 4G’ vs H’). *dlg-RNAi* had very similar effects (Fig. 4E,F).

**Figure 4.**
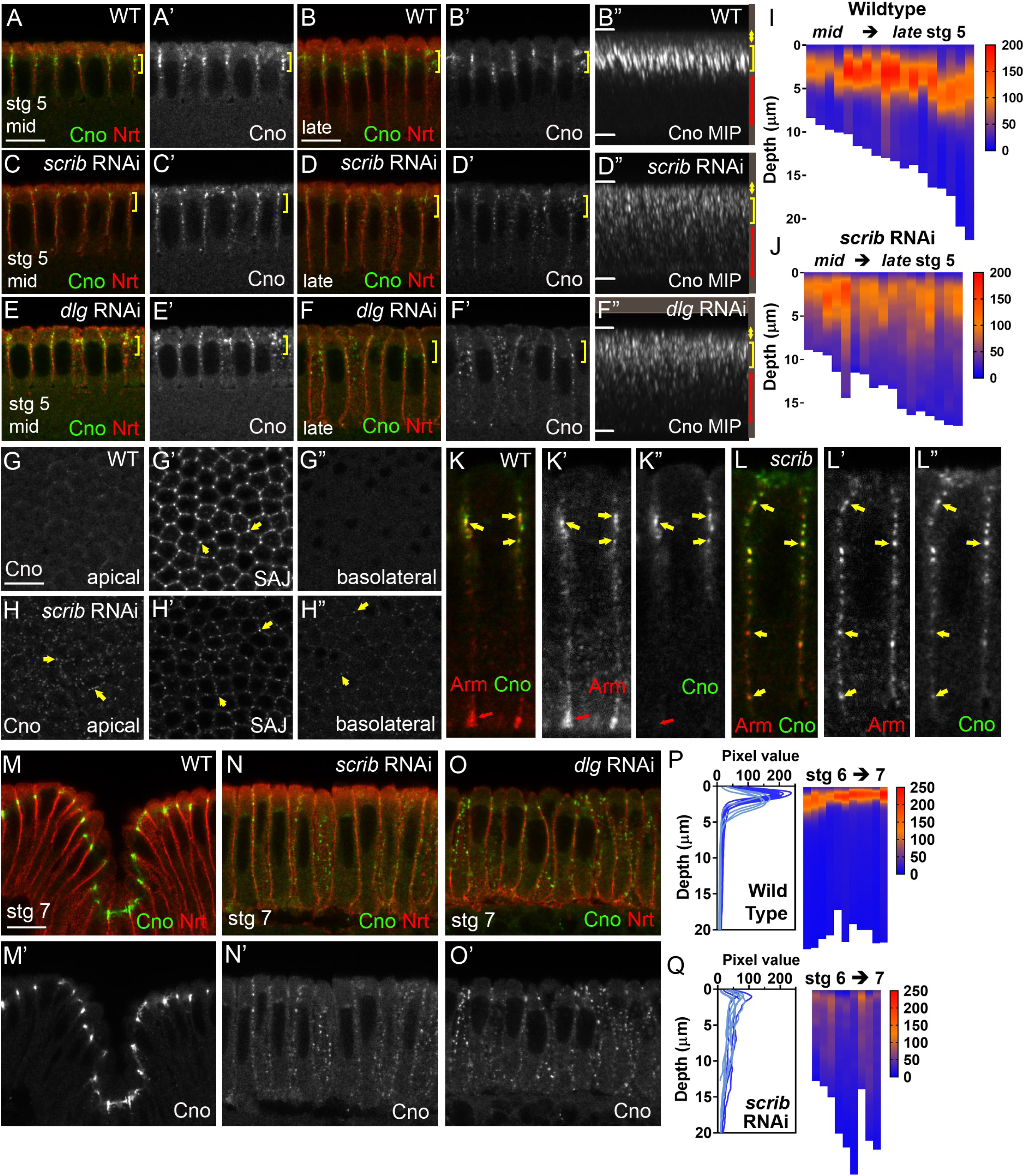
Cno mislocalizes with Arm puncta. A-F,K-O. Cross-sections. B”,D”,F”. MIPs. G,H. *En face* sections through apical membrane (0µm), SAJs (− 3.6µm) and basolateral membrane. I,J,P,Q. Heat maps, Cno puncta displacement along apical-basal axis. Data from MIPs. Each vertical bar=an individual embryo. A-L. Cno enrichment at SAJs (A,B,G,K,M,yellow brackets,arrows) is disrupted after *scrib-RNAi* (C,D,H,L,N) and *dlg-RNAi* (E,F,O). A-H. Cno puncta mislocalize to both apical and basolateral domains during cellularization (stage 5). At SAJ level, Cno is not effectively organized at TCJs (G’,H’). I,J. Corresponding quantification. K,L. Mislocalized Cno puncta track with Arm (yellow arrows). M-O. Cno mislocalization is enhanced during gastrulation (stage 7), with displacement along basolateral domain. P,Q. Corresponding quantification. Scalebars=10µm.

We next asked whether misplaced AJ proteins remained associated or became randomly distributed with respect to one another. In wildtype, Cno and Arm co-localize in apical SAJs (Fig. 4K,yellow arrows), though Cno is more enriched at TCJs and is not included in BJs (Fig. 4K,red arrows). After *scrib-RNAi*, mislocalized Cno and Arm puncta showed clear colocalization along the entire length of the membrane (Fig. 4L,arrows), although Arm’s relative intensity is higher basally. This suggests Scrib or Dlg loss does not affect basic molecular assembly of AJs, but rather supermolecular assembly and retention at the apicolateral boundary.

In wildtype, Cno and AJs move apically at gastrulation onset and tighten into belt AJs (Fig. 4M). Effects of *scrib-RNAi* on Cno became even more pronounced at gastrulation onset. Tight enrichment at apical AJs was completely lost, and instead Cno localized along the apical-basal axis (Fig. 4N, quantified in Fig. 4P vs Q). Some embryos retained residual apical enrichment (Fig. 4Q)—these may be embryos receiving one rather than two copies of the shRNA construct. *dlg-RNAi* similarly disrupted Cno apical enrichment (Fig. 4O). Thus Scrib/Dlg are essential for proper Cno assembly into apical AJs and its retention there during gastrulation.

### Scrib loss reduces Bazooka cortical levels and apical clustering

AJ positioning requires Baz function. In its absence SAJs fail to assemble and cadherin-catenin complexes localize along the lateral membrane. Cno and localized Rap1 activity are required to effectively polarize Baz. Like Cno, Baz localizes in nascent SAJs, and, unlike cadherin-catenin complexes, does not localize further basally (Fig. 5A,brackets). We thus asked if Scrib was essential for directing Baz localization.

**Figure 5.**
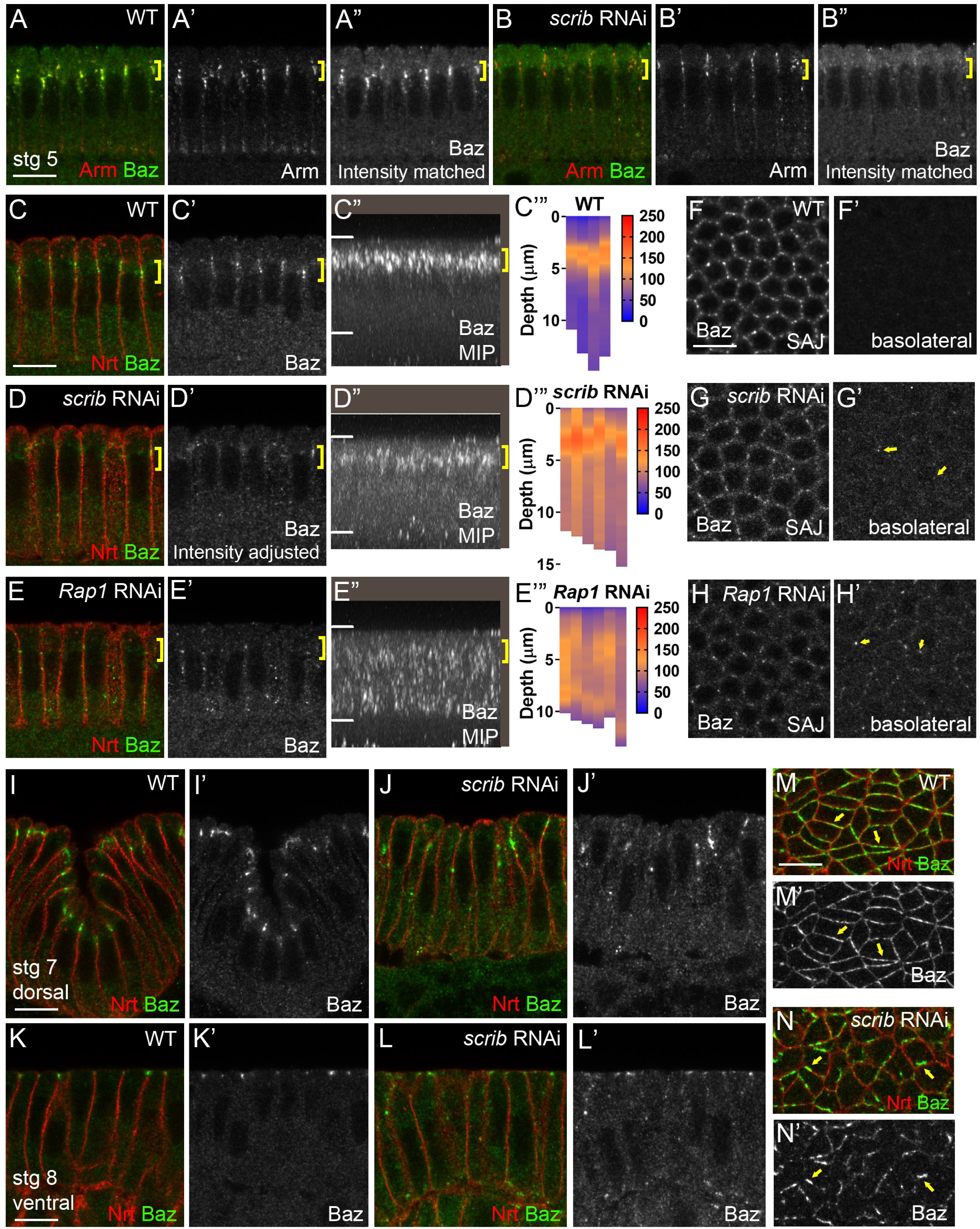
Scrib is required for Baz cortical retention and apical clustering. A,B. Intensity matched images, Arm and Baz in *scrib-RNAi* versus wildtype. *scrib-RNAi* impairs Baz cortical retention during cellularization (yellow brackets). C-E. Intensity-adjusted images, Baz during cellularization. Apical Baz retention is reduced after *scrib-RNAi* (D) compared to complete loss after Rap1 knockdown (E). C”-E”. MIPs and quantification of pixel intensities along apical-basal axis. F-H. *En face* sections. Loss of Scrib or Rap1 both impair Baz clustering at SAJs (− 3.6µm) and mislocalize Baz along the basolateral membrane to varying degrees (yellow arrows). I-L. After gastrulation onset Baz is lost from AJs and displaced as small puncta along the apical-basal axis in *scrib-RNAi*. M,N. *En Face* sections through AJs, stage 7. Circumferential distribution of Baz becomes more irregular. Scalebars=10µm.

*scrib-RNAi* led to reduction in total cortical Baz (Fig. 5A vs B, intensity matched from same experiment). When we elevated the Baz signal to visualize the remaining Baz, we found its tight confinement to SAJs was reduced after *scrib-RNAi* (Fig. 5C vs D; quantified in C’” vs D’”). However, Baz depolarization was more limited than that seen after Rap1 knockdown, in which Cno cortical localization is completely lost (Fig. 5E). Disorganization of Baz in SAJs was also observed *en face* (Fig. 5F vs G,H). The modest pool of Cno retained apicolaterally after *scrib-RNAi* may be sufficient to partially support apical Baz enrichment, albeit at reduced levels and with impaired clustering. Disruption of Baz localization intensified as gastrulation began, with puncta spread along the lateral membrane dorsally (Fig. 3I vs. J) and lacking tight apical localization ventrally (Fig. 5K vs L). Even more striking, the remaining Baz puncta were not uniform around the lateral membrane as in wildtype (Fig. 5M) but formed irregular clusters (Fig. 5N). These were often on dorsal/ventral cell boundaries (Fig. 5N,arrows), potentially reflecting Baz’s normal planar polarization (Fig. 5M,arrows). Thus, Scrib is essential for correctly assembling Baz into nascent SAJs and retaining it in intact junctions as gastrulation begins.

### Scrib/Dlg then maintain junctional and epithelial integrity

In Scrib’s absence epithelial integrity is strongly disrupted (Bilder and Perrimon, 2000), but the time course by which this occurred remained unclear. We thus followed *scrib-RNAi* embryos through gastrulation. In some mutants, including *cno* and *Rap1* (Choi et al., 2013), other mechanisms kick in at gastrulation onset that largely restore apical restriction of AJs and Baz. However, we observed no polarity rescue after Scrib/Dlg knockdown. By wildtype stage 9, Arm, Cno and Baz co-localize in apical AJs (Fig. 6A,C,G,arrows), and only small AJ puncta are seen more basally (Fig. 6D,H). In contrast, after *scrib-RNAi* cells began to separate apically (Fig. 6E,magenta arrow) and AJs were disrupted. Arm and Cno largely colocalized in fragmented apical junctions and more basal puncta (Fig. 6B,E,F,cyan arrows). Baz was even more disrupted, with strong localization limited to larger junctional fragments (Fig. 6I,J,cyan vs magenta arrows). After *scrib-RNAi* cortical actin (Fig. 6K-K”) was altered apically, with F-actin rich protrusions near the apical surface (Fig. 6L,L’), a phenotype also seen in embryos with reduced activity of the formin Diaphanous (Homem and Peifer, 2008) or expressing dominant-negative Rab11 (Roeth et al., 2009). The common phenotype after different perturbations suggest correctly organized AJs restrain actin-driven protrusive behavior.

**Figure 6.**
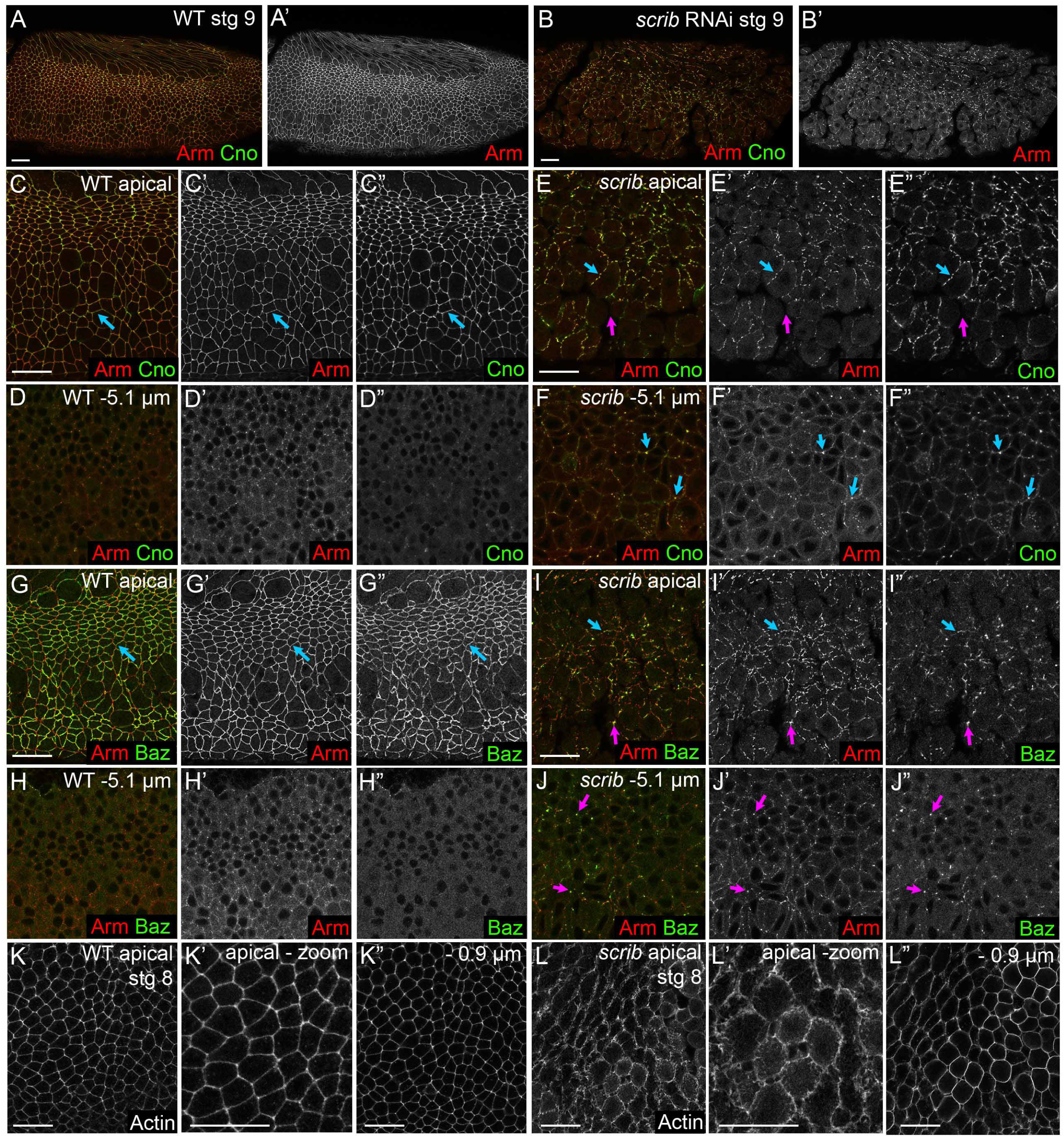
After *scrib-RNAi* AJs fragment and cells lose epithelial character during gastrulation. A-J. By stage 9 *scrib-RNAi* embryos have fragmented AJs (A vs B). In wildtype Arm, Cno, and Baz form continuous belt junctions around cells apically (C,G) with only occasional small Arm puncta more basally (D,H). E,F,I,J. *scrib-RNAi.* Belt AJs fragment (E,I,cyan arrows), gaps form between cells (E,magenta arrow), and junctional fragments appear basally (F,J,arrow). Baz disappears from belt junctions before Arm (I,cyan arrow) and localizes to large junctional fragments (I,J,magenta arrows). K. WT stage 8. Apical actin lines the cortex under AJs (K,K’) and extends along the basolateral cortex (K”). L. *scrib-RNAi.* Apical actin-based protrusions become prominent (L,L’) while basolateral actin is less affected (L”). Scalebars=20µm.

### Scrib/Dlg do not regulate cytoskeletal polarization during cellularization

The cytoskeleton of cellularizing embryos is distinctly polarized (Mavrakis et al., 2009; Schmidt and Grosshans, 2018), reflecting establishment of cytoskeletal polarity during syncytial divisions. Actin is organized into distinct pools: actin-rich microvilli are seen apically, actin lines the basolateral cortex (Suppl. Fig. 3A’) and actin (Suppl.Fig. 3A,arrow, A”) and myosin (Suppl.Fig. 3G,I,K,M,arrows) are strongly enriched at the furrow front. Some probes suggest a pool of actin enrich at apical TCJs (e.g., Sawyer et al., 2009)). Microtubules form inverted baskets over each nucleus (Suppl.Fig. 3D-D”), with nucleation from apical centrosomes. Baz/AJ positioning during cellularization requires both an apical actin scaffold and dynein-mediated retrograde transport (Harris and Peifer, 2005). One mechanism by which Scrib/Dlg knockdown could affect Baz and AJ localization is by disrupting this polarized cytoskeletal organization. However after *scrib-RNAi* or *dlg-RNAi*, F-actin (Suppl.Fig. 3B,C), myosin (Suppl.Fig. 3H,J,L,N), and alpha-tubulin (Suppl.Fig. 3E,F) localization appeared unperturbed. Thus Scrib/Dlg do not act primarily by regulating overall cytoskeletal organization.

### Scrib/Dlg regulates localization of the basolateral polarity protein Par-1, but not all their actions involve Par-1

Par-1 is a basolateral polarity protein that helps restrict localization of junctional and apical proteins. Phosphorylation by Par-1 excludes Baz from the basolateral domain, and manipulating the balance of aPKC and Par-1 activity can re-position AJs (Bayraktar et al., 2006; Benton and St Johnston, 2003; Wang et al., 2012). However, effects of Par-1 loss on Baz localization during polarity establishment are relatively mild, due to functional redundancy with dynein-based apical transport (McKinley and Harris, 2012). We first validated the Par-1 antibody and *par-*1 RNAi tool by immunofluorescence (Suppl.Fig. 4A vs B, C vs D).

In wildtype Par-1 localization changes from cellularization through gastrulation. During cellularization, Par-1 extends along the membrane (Fig. 7A), with highest cortical levels basally (Fig. 7B3) and gradually decreasing levels and less tight cortical localization as it extends apically to overlap SAJs and beyond (Fig. 7B2,B1). Par-1 is gradually cleared from the apical domain during early gastrulation, concentrating basolateral to AJs (Suppl.Fig. 4A, A close-up, E). We next asked whether Scrib/Dlg regulate Par-1 localization during cellularization. After *dlg-RNAi* cortical Par-1 levels were strongly reduced but not eliminated (Fig. 7A vs C). Interestingly, the strongest reduction of Par-1 cortical signal was in the basolateral region (Fig. 7B3 vs D3) rather than near the SAJs (Fig. 7B1 vs D1). *scrib-RNAi* similarly reduced cortical Par-1 (Fig. 7E vs F).

**Figure 7.**
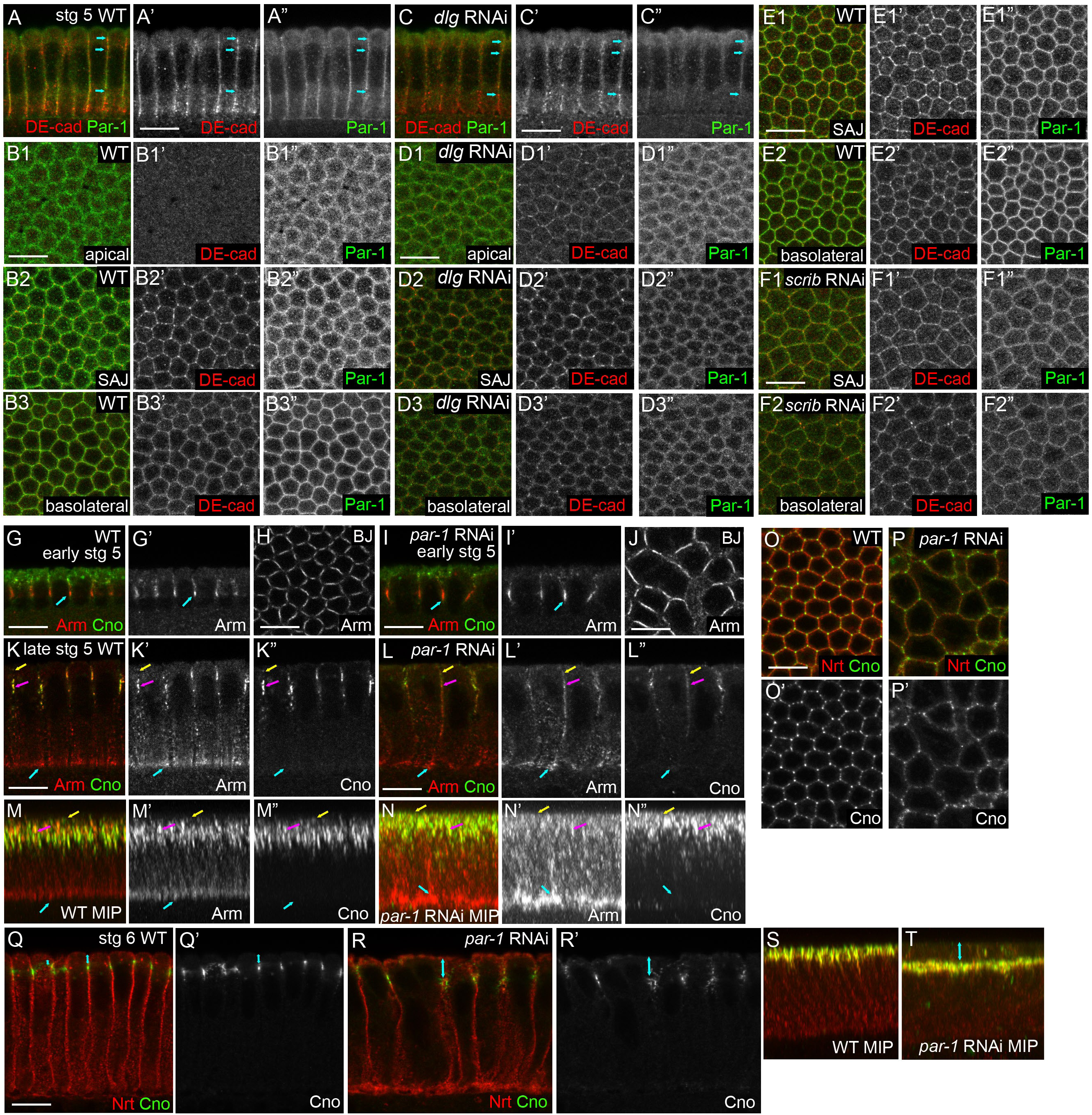
Scrib/Dlg regulate Par-1 localization but Par-1 loss only partially mimics their phenotype. A,C,G,I,K,L,Q,R. Cross-sections. M,N,S,T. MIPs. B,D,E,F,H,J,O,P. *en face* sections at indicated position. A-F. In wildtype (A,B,E) Par-1 localizes in a gradient at the cortex with highest levels basolaterally. Arrows in A indicate positions of sections in B. After *dlg-RNAi* (C,D) or *scrib*-*RNAi* (F) cortical Par-1 is reduced. G-P. Par-1 knockdown reduces Arm enrichment in SAJs during cellularization (K,M vs L,N,magenta arrows) but does not eliminate Arm enrichment in BJs (G,K,M vs I,L,N,cyan arrows, H vs J). Cno is less affected, though it expands apically (K”,M” vs L”,N”,yellow arrows) and is more disorganized in SAJs (O,P). Q-T. As gastrulation begins Arm and Cno refocus into belt AJs in Par-1 knockdown embryos but junctions are positioned more basally (Q vs R, S vs T,arrows). Scalebars=10µm.

This prompted us to examine whether Scrib/Dlg regulate AJ positioning via Par-1, by asking whether *par-1-RNAi* recapitulated Scrib/Dlg knockdown, using a previously characterized shRNA (McKinley and Harris, 2012). *par-1-RNAi* led to strong defects in blastoderm integrity due to partial loss of furrows in syncytial embryos, as previously reported (McKinley and Harris, 2012). This alone suggests Scrib/Dlg knockdown does not completely disable Par-1, as we do not observe this after Scrib/Dlg knockdown. We next examined effects of *par-1-RNAi* on AJs. While *par-1-RNAi* altered polarity establishment, there were several distinctions between Par-1 versus Scrib/Dlg knockdown. First, BJs were largely unaffected by *par-1-RNAi* (Fig. 7G,K,M vs I,L,N,cyan arrows, H vs J). However, SAJs failed to organize appropriately, with the normally tight localization of Arm lost (Fig 7K’ vs L’, M’ vs N’,magenta arrows). Cno and Arm puncta also mislocalized up into the apical domain (Fig 7K’ vs L’, M’ vs N’,yellow arrows). The regular SAJ organization seen *en face* was also lost (Fig. 7O vs P), but this may reflect in part general disorganization of the apical membrane. In strong contrast with Scrib/Dlg knockdown, after *par-1-RNAi* there was partial rescue of AJ localization at gastrulation onset, with Arm and Cno cleared from the basolateral domain and AJs tightening (Fig. 7Q vs R). This is consistent with Baz repolarization observed at this stage (McKinley and Harris, 2012). However, AJs in the dorsal and lateral ectoderm remained abnormally shifted basally (Fig. 7Q vs R, S vs T), as was previously observed (Wang et al., 2012). Thus, Scrib/Dlg are required for Par-1 localization, and Par-1 regulation may account in part for the effect of their loss. However, distinctions in phenotype suggest Par-1 regulation does not account for the full suite of polarity defects observed after Scrib/Dlg knockdown.

### Rab5-dependent trafficking is not essential for Cno junctional clustering but precisely positions AJs

Dynein-dependent trafficking maintains AJ/Baz positioning (Harris and Peifer, 2005), and during later stages DE-cad can traffic through early endosomes (Roeth et al., 2009). Scrib regulates protein trafficking in imaginal discs and other contexts. We thus examined the hypothesis Scrib acts by trafficking DE-cad. RNAi-mediated knockdown of Arm disrupted DE-cad accumulation in SAJs (Harris and Peifer, 2004; Fig 2), and it relocalized to small basolateral puncta and to an apical compartment above nuclei. We suspected this might be an early or recycling endosome. To address this, we localized the early endosome marker Rab5 during cellularization. In wildtype, Rab5 localization was highly reminiscent of mislocalized DE-cad. Rab5 was enriched in an apical compartment above nuclei (Fig. 8A,C,white arrows), in small puncta laterally, and accumulated at the cellularization front (Fig. 8A,C,blue arrows). Rab5 enrichment just above nuclei was particularly clear *en face* (Fig. 8A”,C”). In wildtype apical Rab5 became progressively weaker during gastrulation (Fig. 8E).

**Figure 8.**
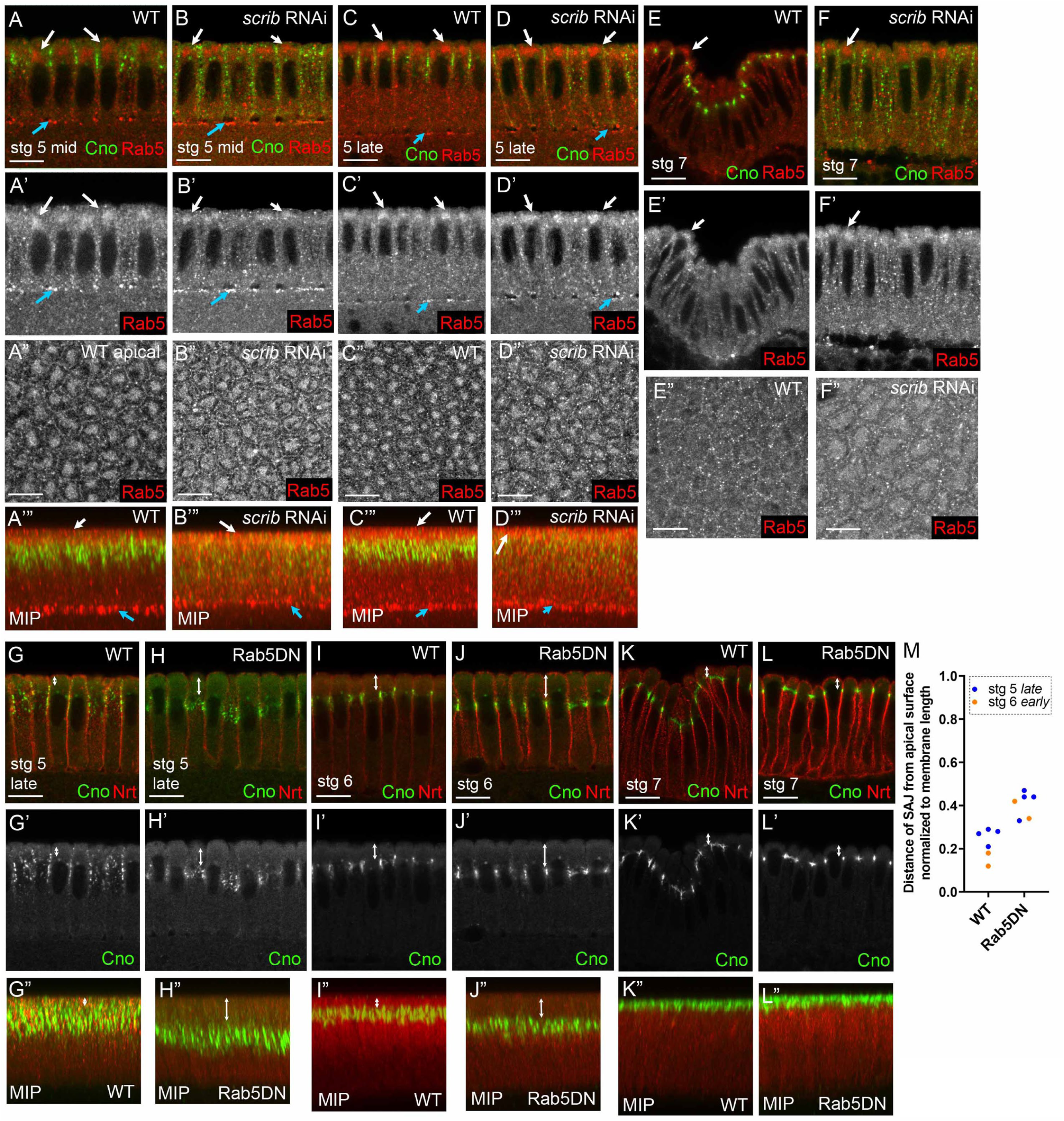
Rab5 localizes to an apical vesicular structure near nascent AJs and regulates precise apical positioning of SAJs. Cross-sections (A-F,G-L), MIPs (A’”-D’”,G”-L”), or *en face* sections at AJ level (A”-F”). A-D. During cellularization of both wildtype and *scrib-RNAi* embryos, Rab5 localizes to the furrow front (cyan arrows) and to an apical vesicular compartment (A-D, A”-D”,A’”-D’”,white arrows). E,F. This apical localization is reduced as gastrulation begins. G-J. Rab5DN does not block Cno assembly into SAJs but does lead to a basal shift in SAJ positioning during cellularization (G vs H,white arrows) and early gastrulation (I vs J,white arrows). K,L. Junctional positioning is restored by stage 7 (K vs L,white arrows). M. Quantification, junctional positioning. Scalebars=10µm.

We thus examined whether *scrib-RNAi* affected Rab5 localization. In *scrib-RNAi* embryos the Rab5 endocytic compartment remained apically polarized (Fig. 8B,D,white arrows), though apical enrichment was mildly reduced (Fig. 8A” vs B”, C” vs D”). To directly test the hypothesis that Scrib/Dlg regulate AJ localization and organization through Rab5-dependent function, we expressed dominant-negative Rab5 (Rab5DN; Zhang et al., 2007). Embryos maternally expressing Rab5DN completed cellularization normally and Cno remained tightly localized and correctly clustered in SAJs. However, SAJs were shift downward along the apical-basal axis--mispositioning continued into early gastrulation (Fig. 8G vs H, I vs J; quantified in M). This effect gradually disappeared as gastrulation proceeded, with Cno focusing apically normally (Fig. 8K vs L, K” MIP vs L” MIP). This unexpected finding is consistent with the idea that SAJ apical-basal polarization is regulated by at least two regulatory inputs – one determining precise SAJ localization along the apical-basal axis and others controlling supermolecular organization. It also suggests Scrib/Dlg do not act by promoting Rab5-regulated AJ trafficking.

### Scrib’s Leucine-rich repeats are essential in polarity establishment and maintenance

Scrib and Dlg are both multi-domain scaffolding proteins with diverse sets of binding partners (Stephens et al., 2018). The overlap between Scrib/Dlg and SAJs is potentially consistent with a simple hypothesis –Scrib and/or Dlg act as direct scaffolds for AJ proteins during polarity establishment. Scrib contains an N-terminal Leucine-rich repeat (LRR) and four C-terminal PDZ domains. Binding partners have been identified for each domain, and mutational analyses have begun to assess their individual roles. In *Drosophila*, *C. elegans* and mammals the LRRs are required for cortical localization and are essential for most known functions (reviewed in Bonello and Peifer, 2019). To examine LRR function, we used a particularly useful missense mutant in Scrib’s LRRs. In *scrib^1^* leucine 223 in the 10^th^ LRR is changed to glutamine (Zeitler et al., 2004). This residue is conserved in all LAP proteins, and structural predictions suggest it is on the surface of the horseshoe-shaped LRR. Expression of a transgenic version in *Drosophila* suggest it encodes a stable protein, though it does not localize to the cortex. *scrib^1^* shares with null alleles loss of polarity in follicle cells, defects in polarity maintenance and epithelial integrity in embryos and loss of polarity and growth regulation in imaginal discs, though transheterozygous interactions with weak alleles suggest it retains a small amount of function (Bilder et al., 2000; Bilder and Perrimon, 2000; Zeitler et al., 2004).

We thus generated females with germlines mutant for *scrib^1^* and crossed them to *scrib^1^* heterozygous males—50% of the progeny are maternally and zygotically mutant while 50% get a wildtype zygotic copy. The cuticle phenotypes matched earlier work—∼50% had the “scribbled” cuticle phenotype, while ∼50% had intact cuticle with morphogenesis defects (Suppl.Fig. 2D,E). We then examined initial polarity establishment. *scrib^1^* mutants recapitulated phenotypes of *scrib-RNAi*. AJ protein localization was completely disrupted in both SAJs (Fig. 9A,B vs E,F,cyan arrows) and BJs (Fig. 9A’,B vs E’,F,magenta arrows), and the normally tight Cno (Fig. 9A”,B’ vs E”, F’,cyan vs magenta arrows) and Baz localization to SAJs (Fig. 9C,D vs G,H,cyan vs magenta arrows) was reduced with spreading basolaterally. Thus, Scrib’s LRRs are essential for its function in polarity establishment. As embryos began gastrulation, polarity disruption was enhanced, with Cno and Arm moving together basolaterally at stage 7 (Fig. 9I vs K), and Baz puncta losing apical localization (Fig. 9J vs L). By stage 9 AJs were fragmented in *scrib^1^* mutants (Fig. 9M vs N). As we observed after *scrib-RNAi*, Arm and Cno co-localized in junctional fragments (Fig. 9O vs Q,arrows), while Baz was restricted to brighter junctional fragments (Fig. 9P vs R,cyan vs magenta arrows). Thus mutating this critical residue in the LRRs essentially eliminated Scrib activity during cellularization and gastrulation.

**Figure 9.**
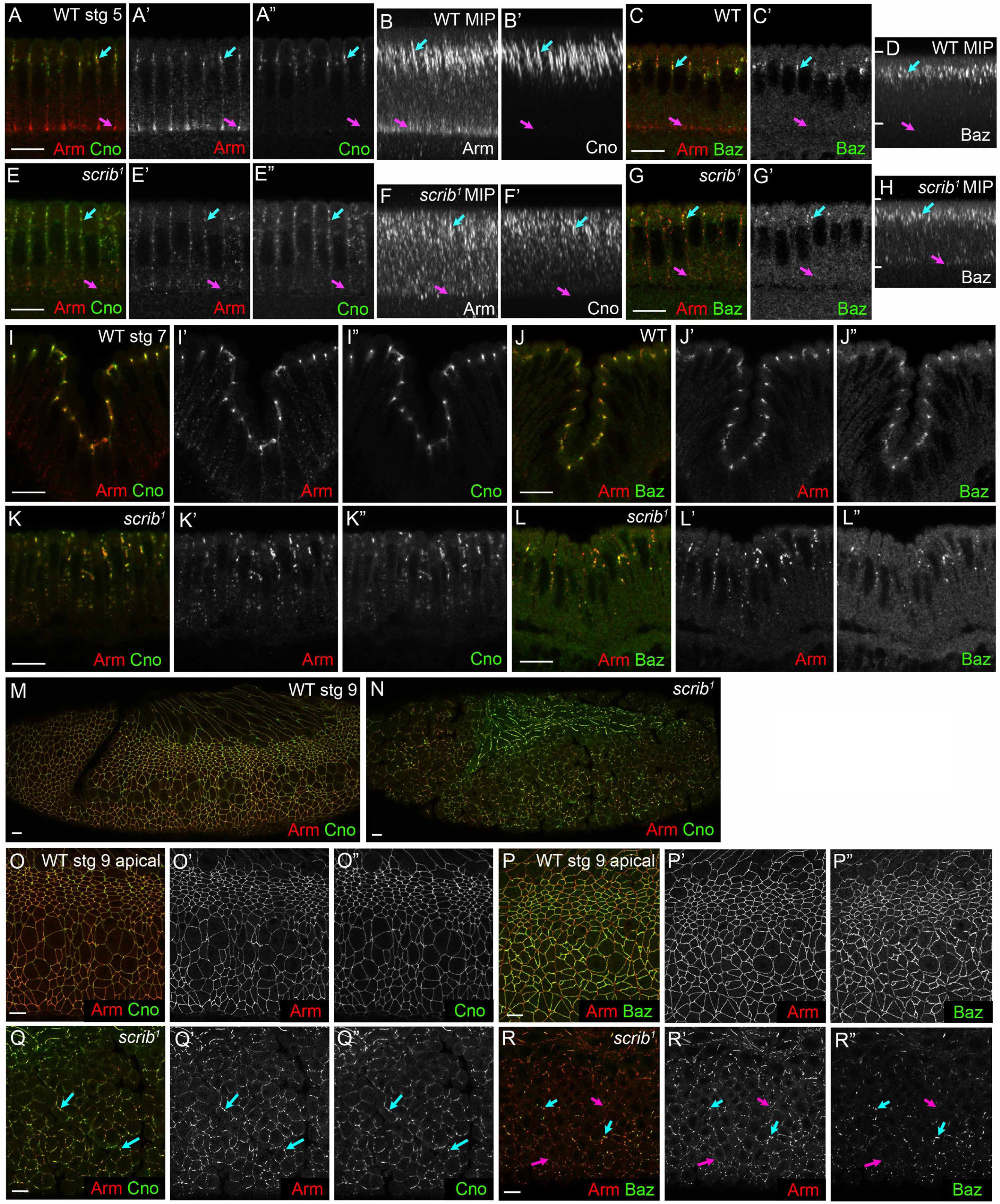
Scrib’s LRR domain is essential for polarity establishment. A-H. Cellularization. Arm enrichment in apical SAJs and BJs (A’,B,cyan and magenta arrows, respectively) is lost in *scrib^1^* mutants (E’,F). Cno positioning and assembly into cables at SAJs (A”,B’) is disrupted in *scrib^1^* mutants, though some apical enrichment remains (E”,F’). The tight apical restriction of Baz in wildtype (C,D) is relaxed in *scrib^1^* mutants (G,H). I-L. Gastrulation onset. Tight apical focusing of Arm, Cno and Baz into belt junctions (I,J) fails in *scrib^1^* mutants, with junctional puncta distributed along the basolateral cortex (K,L). M-R. Stage 9. In wildtype, Arm, Cno and Baz localize continuously in belt AJs (M,O,P). In contrast, in *scrib^1^* mutants AJs fragment and cells begin to separate. Arm and Cno largely co-localize in junctional fragments (Q,arrows), but Baz is more rapidly lost than Arm (R,magenta vs cyan arrows). Scalebars=10µm.

### Scrib’s PDZ domains required to organize SAJs during polarity establishment, but become dispensable for AJs after gastrulation

Intriguingly, while LRR mutations largely or completely inactivate Scrib, the PDZs are not essential for all functions. *scrib^4^*, encoding a truncated Scrib protein lacking all four PDZs, was very informative. This allele rescues apical-basal polarity in embryos and imaginal discs, though it does not rescue SJ assembly or fully rescue growth regulation in imaginal discs (Zeitler et al., 2004). Its embryonic cuticle phenotype illustrates this—while the cuticle is reduced to fragments in *scrib* null mutants, it remains intact in *scrib^4^* mutants, although head involution and dorsal closure are disrupted (Suppl.Fig. 2F).

*scrib^4^* allowed us to selectively disrupt a different subset of Scrib’s protein interactions and examine effects. Consistent with previous reports, embryos maternal/zygotic for *scrib^4^* retained significant ectodermal integrity at late stages (Suppl.Fig 2G vs H; Zeitler et al., 2004)). We next looked during cellularization, to examine whether polarity establishment proceeded normally. To our surprise, *scrib^4^* embryos has defects in SAJs during cellularization. In wildtype, Arm is enriched in apical SAJs and BJs (Fig. 10A,B,cyan and magenta arrows, respectively), while Cno (Fig. 10A”,B’) and Baz (Fig. 10C’,D) are restricted to SAJs. In *scrib^4^* both Arm and Cno localization expanded—they extended apically and basolaterally (Fig. 10A vs E, B vs, F; Arm quantified in 10J), though Cno was still absent from the basal half of the cell. Arm enrichment at basal junctions was also lost in *scrib^4^* (Fig. 10A,B vs E,F,magenta arrows). Baz was less severely affected, with Baz still enriched at the SAJ level (Fig, 10G’,H,cyan arrows), but with Baz puncta also seen more basally. These data are consistent with a role for Scrib’s PDZ domains in initial polarity establishment, but the defects were less severe than those of *scrib^1^* (Fig. 10I).

**Figure 10.**
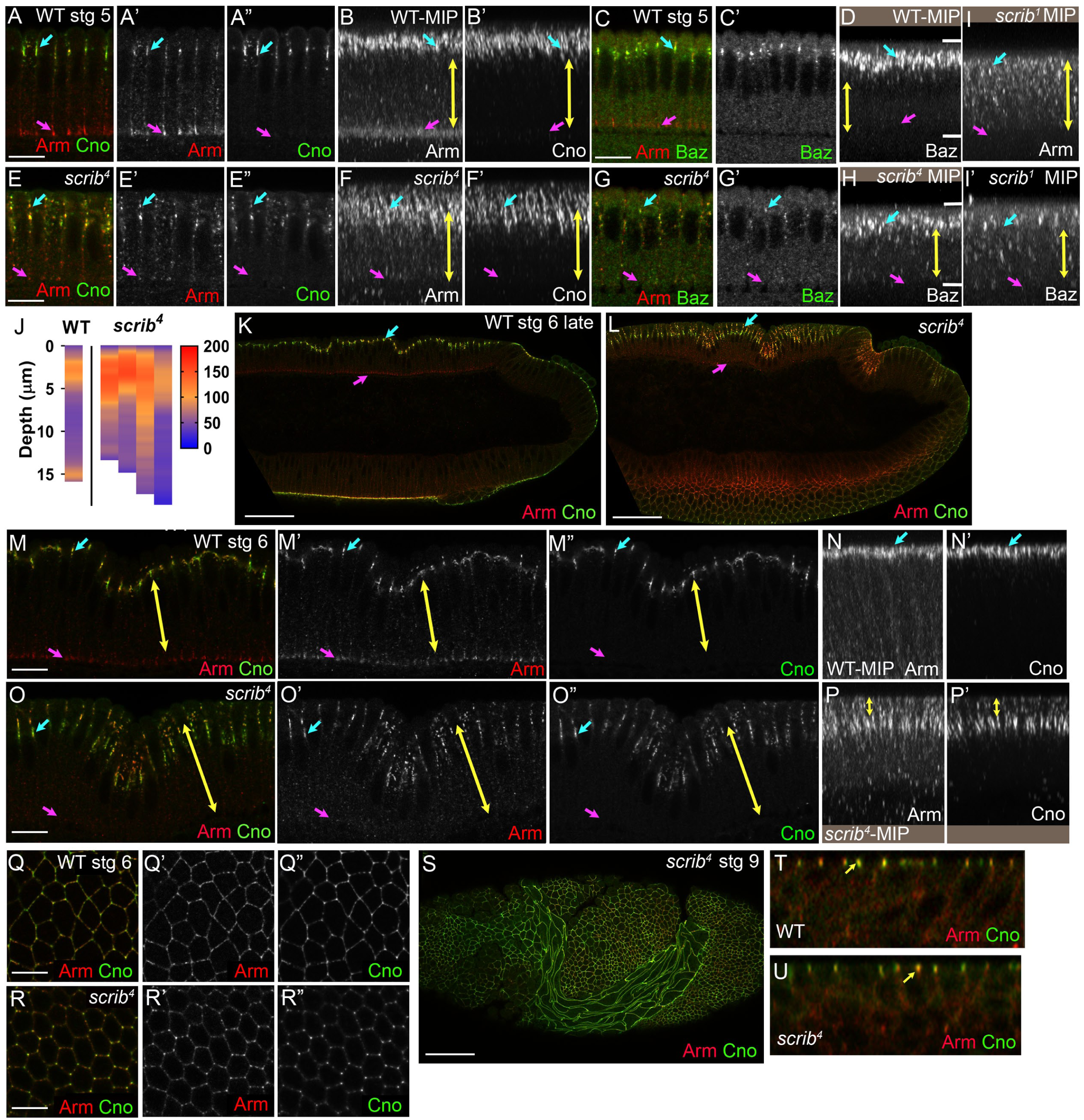
Scrib’s PDZ domains are important for polarity during cellularization but not after gastrulation. A,C,E,G,K,L,M,O. Cross-sections. B,D,F,H,I,N,P. MIPs. Q,R,S. *en face views*, AJ level. A-J. In *scrib^4^* mutants, tight apical localization of Arm and Cno to SAJs during cellularization is reduced (A,B vs E,F,cyan arrows) and Arm enrichment in BJs is lost (magenta arrows). Apical Baz localization broadens (C,D vs G,H,cyan arrows). I. *scrib^1^* effects are more severe. J. Quantification of Arm displacement. K-R. In wildtype Arm and Cno tighten apically both dorsally (K,M,cyan arrows) and laterally (N,cyan arrows). Arm and Cno distribute over a broader region in *scrib^4^* mutants (L,O,P,yellow arrows) with their most intense levels more basal than wildtype (O,cyan arrows). *scrib^4^* mutants are delayed in the SAJ to belt AJ transition (Q vs R). S-U. In stage 9 *scrib^4^* mutants cell shapes remain abnormal (S) but junctional proteins repolarize (T,U,yellow arrows). K,L,S Scalebars=50µm. All others=10µm.

We then followed *scrib^4^* mutants to see when and how polarity was re-established. In wildtype, AJs move apically and become more focused by late stage 6 (Fig. 10K,M,N,cyan arrows), though some Arm remains in BJs (Fig. 10K,M’,magenta arrows). Focusing of Arm and Cno into AJs was delayed in *scrib^4^*—both extended further basolaterally (Fig. 10O,yellow arrows), and in the lateral epidermis their strongest localization was more basal than in wildtype (Fig. 10P,yellow arrows). The transition from SAJs to belt AJs in *scrib^4^* was also delayed (Fig. 10Q vs R). Remarkably, however, AJ repolarization was complete by the end of germband extension (Fig. 10S, z-stack reconstructions T vs U), unlike what we observed after *scrib-RNAi*. Thus Scrib’s PDZ domains are required for many aspects of polarity establishment, but PDZ domain-independent mechanisms appear during gastrulation and restore polarity.

## Discussion

Identifying the earliest symmetry-breaking events that initially position AJs, thereby setting the boundary between apical and basolateral domains, is a key aspect of understanding how polarity is established. Here we report that Scrib/Dlg, best known for roles as basolateral determinants during polarity maintenance, play a separate and surprising role in organizing AJs during polarity establishment, positioning them at the top of the polarity network.

### Master directors of junction supermolecular organization

Scrib and Dlg are multidomain proteins with many partners, allowing them to serve diverse biological functions, from synaptogenesis to oriented cell division. Our data reveal they play distinct roles during polarity establishment and polarity maintenance, likely engaging very different sets of binding partners. This is supported by the evolving localization pattern of Scrib/Dlg on the plasma membrane, with sequential co-localization with and roles in positioning AJ versus SJ proteins, suggesting the capacity to engage with and position distinct junctional and polarity proteins. Our analyses also begin to dissect the underlying molecular basis. Scrib’s PDZ domains are important for the precision of initial polarity establishment but are redundant with other mechanisms for polarity maintenance after gastrulation, though they regulate SJ positioning.

AJs play a key role at the boundary between apical and basolateral domains, and building a functional junction is a multistep process. This includes assembling the core cadherin-catenin complex, positioning it, and supermolecular assembly. Assembly of the core complex appears to occur coincident with synthesis, and thus small puncta are already present before cellularization. As cellularization proceeds, these are captured at the apicolateral interface in a process requiring Baz, Cno, and an intact actin cytoskeleton, where they coalesce into SAJs, with ∼1500 AJ complexes and 200 Baz proteins. Cadherin-catenin complexes form independently of either Baz or Cno, but AJ positioning and full supermolecular assembly depend on both. We found Scrib/Dlg are also key for AJ apicolateral retention and supermolecular assembly, although Arm and Cno remain associated in misplaced puncta, and thus core AJ complexes remain intact. Further, a second junctional complex that arise during polarity establishment, the BJs, also require Scrib/Dlg for its supermolecular organization. Unlike AJs, BJ organization is not dependent on other polarity determinants including Cno, Rap1 or Par-1. It will be of interest to examine if Scrib/Dlg act via known regulators of cadherin clustering, including intrinsic (e.g. cis- and trans-interactions of cadherins) and extrinsic (e.g. local actin regulation, endocytosis) factors (Truong Quang *et al*., 2013).

### The Scrib/Dlg module shapes polarity establishment and maintenance via multiple mechanisms

Our ultimate goal is to define molecular mechanisms underlying polarity establishment. Our new data place Scrib/Dlg in a critical position near the top of the network, but also suggest they act via multiple effectors. Perhaps the strongest evidence for multiple roles with distinct effectors comes from analysis of *scrib^4^*. Supermolecular organization of both SAJs and BJ must involve interactions with specific partners via the PDZ domains— one speculative possibility is that these include core AJ proteins, as ßcatenin can coIP with Scribble and interact with PDZ domains 1 and 4 (Ivarsson et al., 2014; Zhang et al., 2006). Testing this idea will be an important future direction. This initial role may also involve modulating Par-1. During cellularization, Scrib/Dlg and Par-1 localize in “inverse gradients”: Scrib and Dlg enriched at the SAJ level, and Par-1 with higher cortical intensity basolaterally. Scrib/Dlg play a role in effective membrane recruitment of Par-1 at this stage, and effects of *par-1-RNAi* on SAJ protein localization during cellularization are largely similar to those of *scrib-RNAi*. However, regulating Par-1 is not the only mechanism by which Scrib/Dlg act, as AJs are rescued during gastrulation after *par-1-RNAi*.

Scrib then plays a second PDZ-independent role as gastrulation begins, ensuring focusing of cadherin-catenin complexes and Baz into apical belt AJs. This requires the N-terminal LRRs but not the PDZs. Positioning Baz at this stage involves at least two inputs which are redundant with one another, one via Par-1 and one via an apical transport mechanism (McKinley and Harris, 2012). One speculative possibility is that Scrib/Dlg also regulate protein trafficking, a role they have in other contexts. However, disrupting Scrib/Dlg function has very different consequences than disrupting Rab5-dependent trafficking, suggesting they do not act via Rab5. aPKC also provides important cues at this stage—perhaps Scrib/Dlg regulate aPKC localization or function. It will be important to further explore the nature of this second role.

### Defining the ultimate upstream cue for polarity establishment

Our initial goal more than a decade ago was to define roles of AJs in polarity establishment. However, it rapidly became apparent AJs are not at the top of the hierarchy. Cno, Rap1 and Baz act upstream of AJ positioning and supermolecular assembly. Our new data moves us another step upward in the network, revealing a key role for Scrib/Dlg in regulating AJ positioning and assembly. However, they also reveal that the process is not a simple linear pathway, and raise new questions. Loss of Scrib or Dlg almost completely disrupts AJs during cellularization. However, effects on Baz and Cno are less complete—supermolecular assembly is affected, but they are retained in the apical half of the membrane. This suggests other cues are involved. The ultimate polarizing cue during syncytial development is the oocyte membrane, which then directs cytoskeletal polarization. Cytoskeletal cues regulate Cno localization. While our data rule out a role for Scrib/Dlg in establishing basic cytoskeletal polarity, they do not rule out roles, for example, in localizing a special “type” of actin cytoskeleton in the apical domain. Retention of Cno at the membrane after Scrib/Dlg knockdown suggests that minimally basal Rap1 activity remains intact. Changes to early Par-1, and to a lesser extent Baz, cortical localization with loss of Scrib/Dlg, also raise the possibility that lipid-based regulation is impaired (Kullmann and Krahn, 2018; McKinley et al., 2012). At this time, we do not know what cues regulate Scrib/Dlg apical enrichment but AJs do not appear to direct this, nor are they essential for polarizing Cno or Baz. Continued characterization of the full protein network and molecular mechanisms governing polarity establishment will keep the field busy for years to come.

## Materials and Methods

**Table 1:** Fly stocks, antibodies, and probes.

| Fly Stock |  | Source |
| --- | --- | --- |
| UAS.*scrib* RNAi (valium20; BL35748) |  | Bloomington *Drosophila* Stock Center (BDSC) |
| UAS.*scrib* RNAi (valium22; BL38199) |  | BDSC |
| UAS.*dlg1* RNAi (valium22; BL35286) |  | BDSC |
| UAS.*arm* RNAi (valium20; BL35004) |  | BDSC |
| UAS.*Rap1* RNAi (valium20; BL35047) |  | BDSC |
| UAS.*par-1* RNAi (valium20; BL32410) |  | BDSC |
| UASp.YFP.Rab5.S43N (BL9771) |  | BDSC |
| FRT82B ovoD1-18 (BL2149) |  | BDSC |
| hsFLP;;Dr[Mio]/Tm3 (BL7) |  | BDSC |
| *scrib*<sup>1</sup> FRT82B/Tm3 |  | Kindly provided by David Bilder (Bilder and rimon, 2000) |
| *scrib*<sup>4</sup> FRT82B/Tm6C |  | Kindly provided by David Bilder (Zeitler et al., 2004) |
| Antibodies/probes | Dilution (IF, WB*) | Source |
| **Primary antibodies** |  |  |
| Anti-Nrt (mouse) | 1:100 | BP106, Developmental Studies Hybridoma Bank (DSHB) |
| Anti-Arm (mouse) | 1:100 | N27A1, DSHB |
| Anti-DE-cad (rat) | 1:100 | DCAD2, DSHB |
| Anti-Dlg (mouse) | 1:500, 1:1000* | 4F3, DSHB |
| Anti-Cno (rabbit) | 1:1000 | J. Sawyer and N. Harris (University of North Carolina at Chapel Hill) |
| Anti-Baz (rabbit) | 1:2000 | (Choi et al., 2013) |
| Anti-Scrib (guinea-pig) | 1:500, 1:1000* | (Zeitler et al., 2004) |
| Anti-Par-1 (guinea-pig) | 1:800 | (McDonald et al., 2008) |
| Anti-Rab5 (guinea-pig) | 1:1000 | (Tanaka and Nakamura, 2008) |
| Anti-Zip (rabbit) | 1:500 | (Chougule et al., 2016) |
| Anti-α-Tubulin (mouse) | 1:4000, 1:5000* | T6199, Sigma Life Science |
| Phalloidin Alexa 488 | 1:500 | A12379, Thermo Fisher Scientific |
| **Secondary antibodies** |  |  |
| Alexa 488, 568, and 647 | 1:500 | Life Technologies |
| IRDye 680DR and RDye 800CW | 1:10000 | Licor Biosciences |

### Fly genetics

Fly stocks used in this study are listed in Table 1. Mutations are described at FlyBase (http://flybase.org). Wild type was yellow white. All experiments were carried out at 25°C, with the exception of driving *arm* RNAi which was done at 27°C. Knockdown by RNAi was achieved by crossing double copy maternal GAL4 females (Staller et al., 2013) to males carrying the UAS hairpin construct. Female progeny were then crossed back to males carrying the UAS hairpin construct. For *scrib* RNAi most experiments were carried out with the valium 22 line, but key conclusions were verified with the valium 20 line. For driving *arm* RNAi expression the GAL4 was provided by the males (Ni et al., 2011). Maternal expression of dominant negative Rab5 was carried out by crossing female double maternal GAL4 flies to males carrying UAS-YFP.Rab5.S43N. To create germline clones, *scrib1* or *scrib4* females were crossed to males carrying hsFLP;; FRT82B ovoD1. Wandering larvae were heat shocked for 2 h in a 37°C water bath on two consecutive days.

### Immunofluorescence

Staining for Nrt, Arm, Cno, Baz, Scrib, Rab5 and Zip were performed using the heat-fix method. Dechorionated embryos were fixed in boiling Triton salt solution (0.03% Triton X-100, 68 mM NaCl) for 10 s followed by fast cooling on ice and devitellized by vigorous shaking in 1:1 heptane:methanol. Embryos were stored in 95% methanol/5% EGTA for at least 48 h at −20°C prior to staining. For staining all other antigens, embryos were fixed for 20 min with 9% formaldehyde and devitellized in 1:1 heptane:methanol, except for staining with phalloidin where embryos were devitellized in 1:1 heptane:90% ethanol. Prior to staining embryos were washed with 0.1% Tween-20 in PBS (heat fixed embryos) or 0.1% Triton X-100 (formaldehyde fixed embryos), and blocked in 1% normal goat serum in PBST for 1 h. Primary and secondary antibody dilutions are listed in Table 1.

### Image acquisition and manipulation

Fixed embryos were mounted in Aqua-poly/Mount and imaged on a confocal laser-scanning microscope (LSM 880; 40x/NA 1.3 Plan-Apochromat oil objective; Carl Zeiss, Jena Germany). Images were processed using ZEN 2009 software. Photoshop CS6 (Adobe) was used to adjust input levels so that the signal spanned the entire output grayscale and to adjust brightness and contrast.

### Image quantification

Maximum Intensity Projections (MIPs) were generated by acquiring z-stacks through the embryo with a 0.3 µm step size and digital zoom of 2. ZEN 2009 software was used to crop stacks to 250×250 pixels along the xy-axis and to project xyz-stacks along the y-axis as previously described (Choi et al., 2013). The apical-basal position of puncta was determined from MIPs using ImageJ. Projections were rotated 90° counterclockwise and analyzed using the Plot Profile function in ImageJ (NIH) to generate values of average fluorescence intensity along the apical-basal axis. These data were displayed as heat maps, illustrating intensity along the apical-basal axis with a color gradient. Graphs and accompanying heat maps were generated using GraphPad Prism 8.0.

### Cuticle preparation

Cuticle preparation was performed according to Wieschaus and Nusslein-Volhard (Wieschaus and Nüsslein-Volhard, 1986).

### Immunoblotting

Knockdown efficiency of *scrib* and *dlg* were determined by western blotting. Embryo lysates were prepared by grinding dechorionated embryos in ice-cold lysis buffer (1% NP-40, 0.5% Na deoxycholate, 0.1% SDS, 50 mM Tris pH8, 300 mM NaCl, 1.0 mM DTT and Halt^TM^ protease and phosphatase inhibitor cocktail [Thermo Scientific]). Lysates were cleared at 16,000 g and protein concentration determined using the Bio-Rad Protein Assay Dye (Bio-Rad). Lysates were resolved by 8% SDS-PAGE, transferred to nitrocellulose filters and blocked for 1 h in 10% dry milk powder in PBS-Tw (0.1% Tween-20 in PBS). Membranes were incubated in primary antibody for 2 h (see Table 1 for antibody concentrations); washed four times for 5 min each in PBS-Tw; and incubated with IRDye-coupled secondary antibodies for 45 min. Signal was detected with the Odyssey infrared imaging system (Licor Biosciences). Band densitometry was performed using ImageStudio software version 4.0.21 (LI-COR).

## Acknowledgements

We thank David Bilder and Mark Khoury for Scribble antibody, *scribble* mutant stocks, advice in generating maternal-zygotic mutants, and helpful discussions Jocelyn McDonald and Jeffrey Thomas, the Bloomington Drosophila Stock Center, the Transgenic RNAi Project (TRiP) and the Developmental Studies Hybridoma Bank for reagents, Tony Perdue of the Biology Imaging Center, Clara Williams, Jonathan Deliberty and Ian Windham for technical assistance, John Poulton for reagents and thoughtful discussion, Scott Williams and Peifer lab members for helpful advice and comments, and Bing He for helpful discussions. This work was supported by NIH R35 GM118096 to M.P. T.T.B. was supported in part by a Sir Keith Murdoch Fellowship from the American Australian Association.

## Competing Interests

No competing interests declared

---

**Supplemental Figure 1. Our *scrib* and *dlg-RNAi* tools effectively knockdown protein levels.** Representative Western blots of embryonic extracts at the indicated ages in wildtype and after expression of our chosen RNAi lines, driven by a pair of strong maternal GAL4 drivers (MatGAL4). A. Two independent *scrib-RNAi* lines effectively knockdown maternal and zygotic Scrib during early development and mid-embryogenesis. Most experiments were carried out with the valium 22 line, but key conclusions were verified with the valium 20 line. B. Effective knockdown of maternal and zygotic Dlg by the *dlg*-*RNAi* line used in this study. C. *scrib-RNAi* did not substantially alter Dlg levels, nor did *dlg-RNAi* alter Scrib levels.

**Supplemental Figure 2. Comparison of cuticle phenotypes between *scrib-RNAi*, *dlg-RNAi* and *scrib* mutants**. A-F. Cuticles, anterior up. A. Wildtype. Note intact head skeleton (black arrow) and intact epidermis with alternating denticle belts (magenta arrow) and naked cuticle. B-D. *scrib-RNAi* (B) and *dlg-RNAi* (C) phenocopy the strong *scrib^1^* allele (D) showing the characteristic “scribbled” cuticle phenotype with remnant vesicles and tubules of cuticle (arrows). E. Paternally rescued *scrib^1^* mutant—morphogenesis, including head involution (black arrow) and dorsal closure (red arrow) fail, but epidermal cuticle integrity is largely restored. F. Maternal/zygotic *scrib^4^* mutant. Head involution (black arrow) and dorsal closure (red arrow) fail. G,H. Wildtype vs. *scrib^4^* mutant during dorsal closure. In the *scrib^4^* mutant the epidermis is intact but head involution has failed (arrow). Scale bars=50µm.

**Supplemental Figure 3. Neither Scrib nor Dlg knockdown alters basic cytoskeletal polarity.** Embryos, mid- to late cellularization, genotypes and antigens indicated. A. In wildtype, actin is found all along the cortex, including at the level of the nascent AJs (A’) but is substantially enriched at the cellularization front and the closing basal rings (A arrow, A”). B,C. This is unaltered after *dlg* or *scrib-RNAi*. D. In wildtype tubulin forms inverted baskets (D,D”) nucleated from centrosomes above each nucleus (D, arrow, D’). E,F. This is unaltered after *dlg* or *scrib-RNAi*. G,I,K,M. Myosin localizes to the cellularization front (arrows) and more diffusely to an apical pool. H,J,L,N. Once again, these features are unaltered after *dlg* or *scrib-RNAi*. Scale bar=20µm.

**Supplemental Figure 4. Par-1 and the AJs largely segregate by stage 8.** Embryos, stage, genotype and antigens indicated. A,C. By stage 7 AJs move apically and localize largely apical to Par-1 (A yellow arrows, closeup, C1), while Par-1 remains on the basolateral cortex (A cyan arrows, C2). B,D. Validation of *par-1*-*RNAi* reagent. *par-1*-*RNAi* essentially eliminated Par-1 signal, as detected by immunofluorescence. E. At stage 8 AJs continue localize apical to Par-1. Scale bar=20µm.

## Notes

#### Summary of Updates

Author order was entered incorrectly

