## Supplemental Figures for "Scribble and Discs-large direct adherens junction positioning and supermolecular assembly to establish apical-basal polarity"

**Supplemental Figure 1. Our *scrib* and *dlg-RNAi* tools effectively knockdown protein levels.** Representative Western blots of embryonic extracts at the indicated ages in wildtype and after expression of our chosen RNAi lines, driven by a pair of strong maternal GAL4 drivers (MatGAL4). A. Two independent *scrib-RNAi* lines effectively knockdown maternal and zygotic Scrib during early development and mid-embryogenesis. Most experiments were carried out with the valium 22 line, but key conclusions were verified with the valium 20 line. B. Effective knockdown of maternal and zygotic Dlg by the *dlg*-RNAi line used in this study. C. *scrib-RNAi* did not substantially alter Dlg levels, nor did *dlg-RNAi* alter Scrib levels.

### Suppl. Figure 1

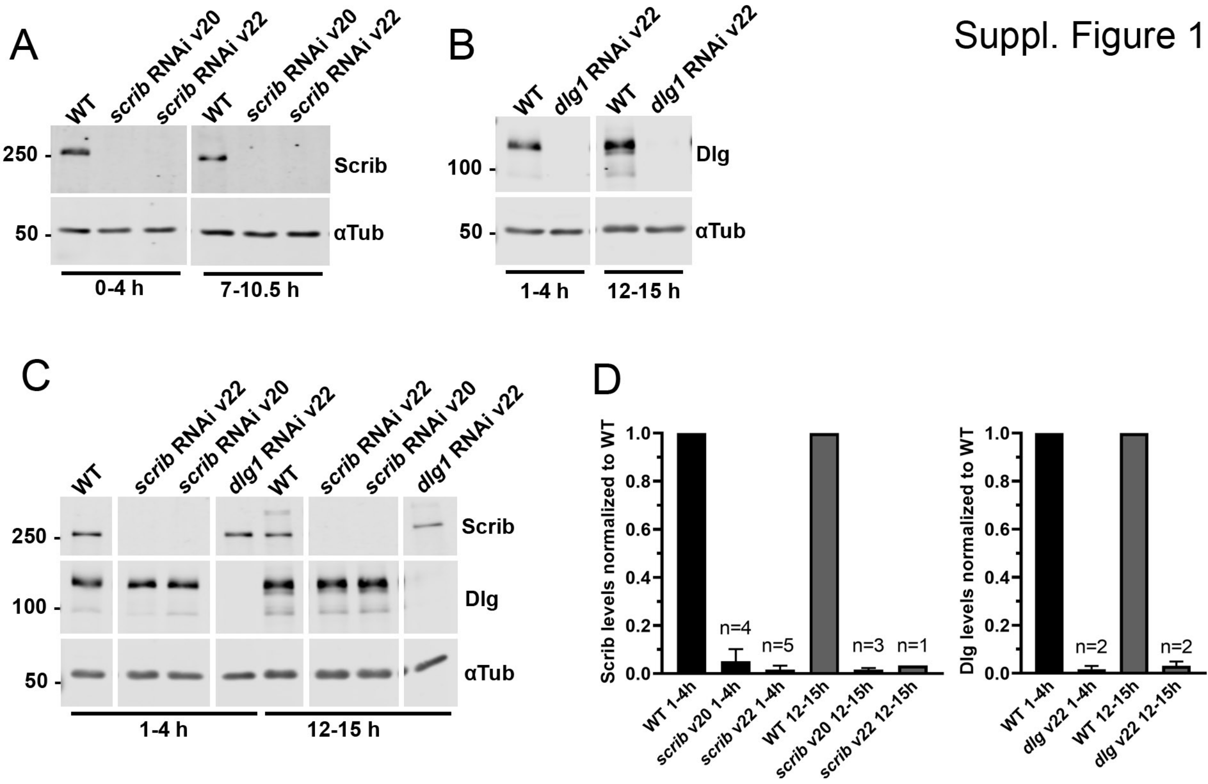

**Supplemental Figure 2. Comparison of cuticle phenotypes between *scrib-RNAi*, *dlg-RNAi* and *scrib* mutants**. A-F. Cuticles, anterior up. A. Wildtype. Note intact head skeleton (black arrow) and intact epidermis with alternating denticle belts (magenta arrow) and naked cuticle. B-D. *scrib-RNAi* (B) and *dlg-RNAi* (C) phenocopy the strong $scrib^1$ allele (D) showing the characteristic "scribbled" cuticle phenotype with remnant vesicles and tubules of cuticle (arrows). E. Paternally rescued $scrib^1$ mutant—morphogenesis, including head involution (black arrow) and dorsal closure (red arrow) fail, but epidermal cuticle integrity is largely restored. F. Maternal/zygotic $scrib^4$ mutant. Head involution (black arrow) and dorsal closure (red arrow) fail. G,H. Wildtype vs. $scrib^4$ mutant during dorsal closure. In the $scrib^4$ mutant the epidermis is intact but head involution has failed (arrow). Scale bars=50μm.

Suppl. Fig 2

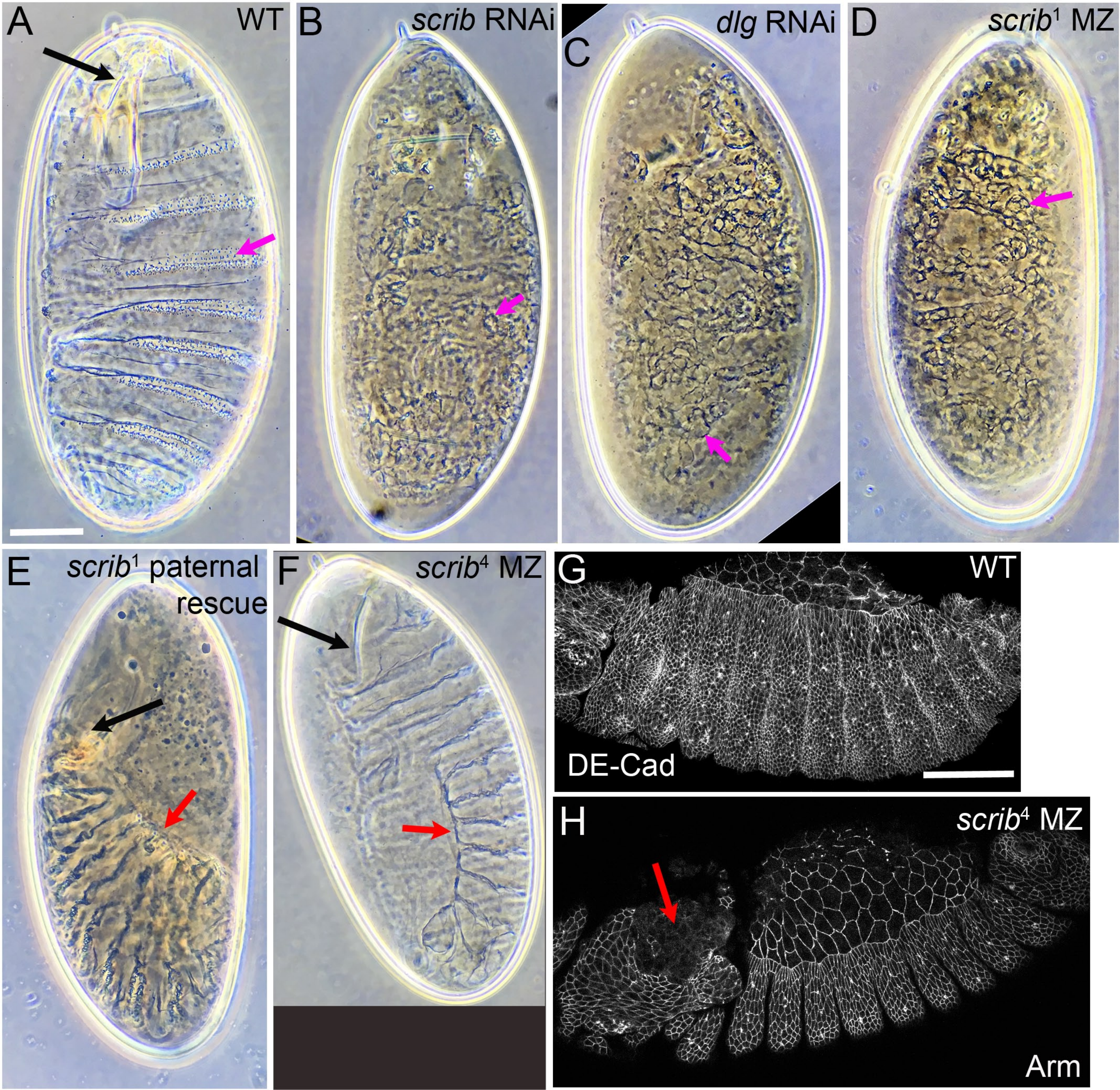
A
WT
B
scrib RNAi
C
dlg RNAi
D
scrib1 MZ
E
scrib1 paternal rescue
F
scrib4 MZ
G
WT
DE-Cad
H
scrib4 MZ
Arm

**Supplemental Figure 3. Neither Scrib nor Dlg knockdown alters basic cytoskeletal polarity.** Embryos, mid- to late cellularization, genotypes and antigens indicated. A. In wildtype, actin is found all along the cortex, including at the level of the nascent AJs (A') but is substantially enriched at the cellularization front and the closing basal rings (A arrow, A"). B,C. This is unaltered after *dlg* or *scrib-RNAi*. D. In wildtype tubulin forms inverted baskets (D,D") nucleated from centrosomes above each nucleus (D, arrow, D'). E,F. This is unaltered after *dlg* or *scrib-RNAi*. G,I,K,M. Myosin localizes to the cellularization front (arrows) and more diffusely to an apical pool. H,J,L,N. Once again, these features are unaltered after *dlg* or *scrib-RNAi*. Scale bar=20μm.

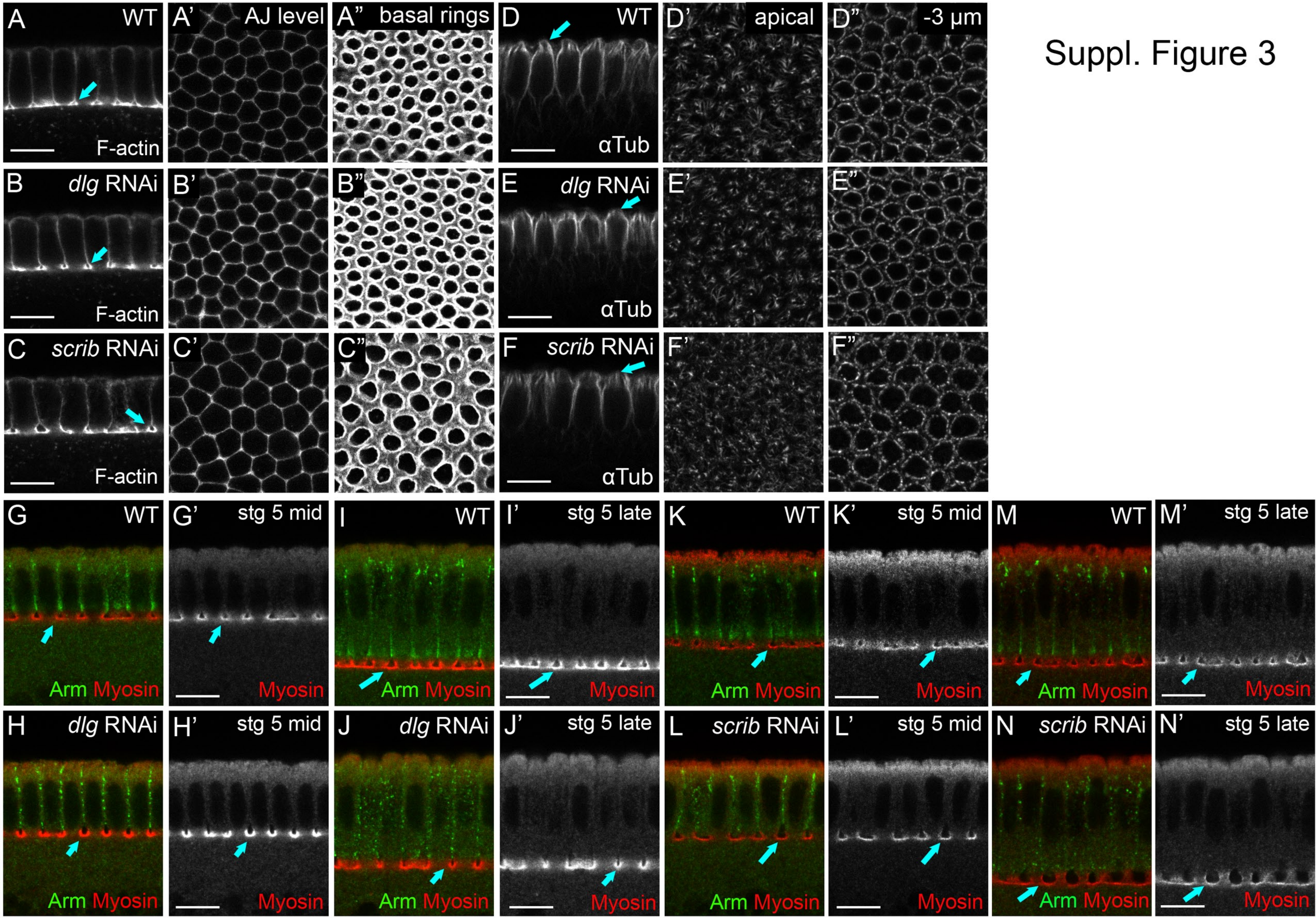


Suppl. Figure 3

**Supplemental Figure 4. Par-1 and the AJs largely segregate by stage 8.** Embryos, stage, genotype and antigens indicated. A,C. By stage 7 AJs move apically and localize largely apical to Par-1 (A yellow arrows, closeup, C1), while Par-1 remains on the basolateral cortex (A cyan arrows, C2). B,D. Validation of *par-1* RNAi regent. *par-1* RNAi essentially eliminated Par-1 signal, as detected by immunofluorescence. E. At stage 8 AJs continue localize apical to Par-1. Scale bar=20μm.

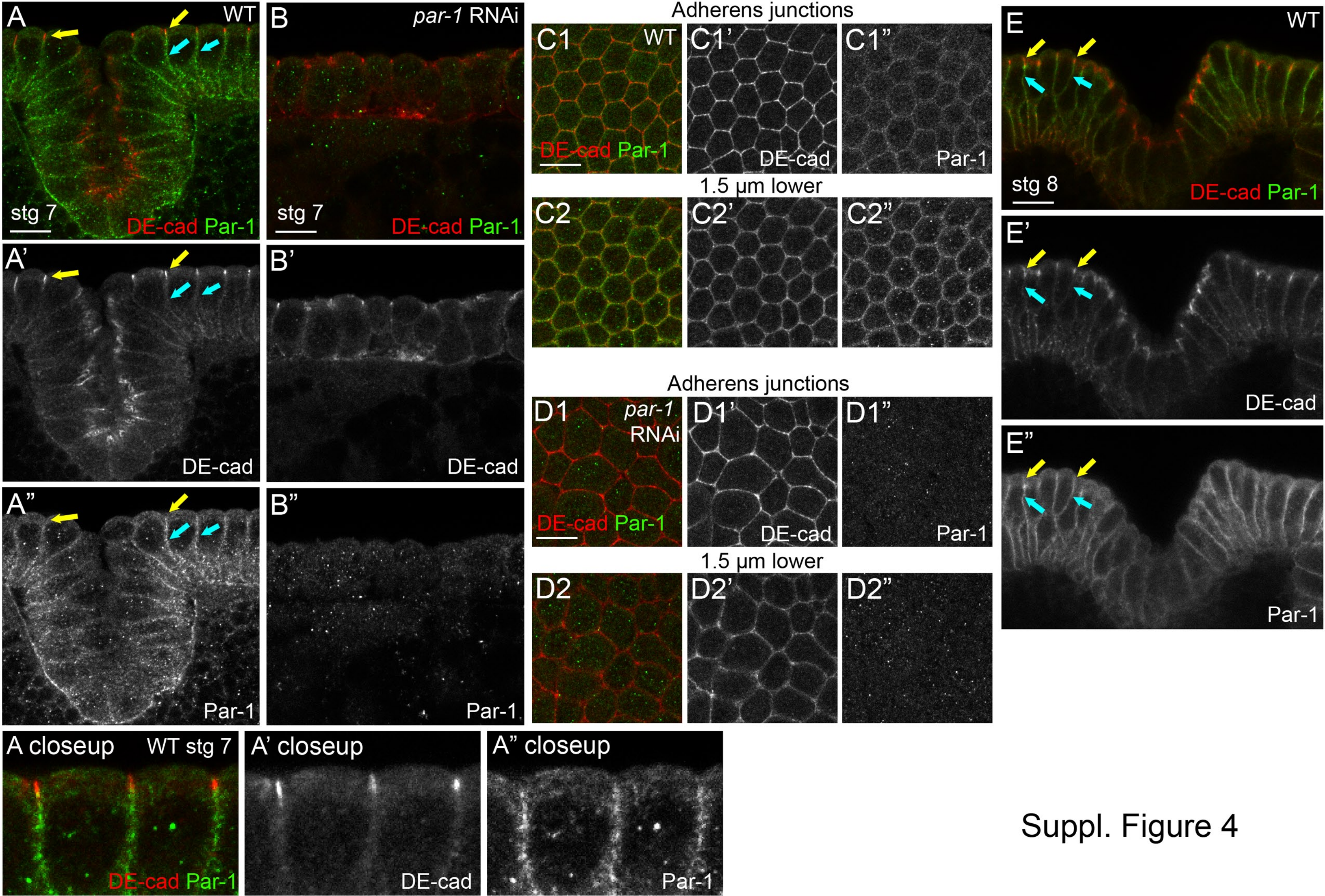


Suppl. Figure 4
